# Angiopoietin-like 4 shapes the intrahepatic T-cell landscape via eIF2α signaling during steatohepatitis in diet-induced NAFLD

**DOI:** 10.1101/2023.01.10.523354

**Authors:** Zun Siong Low, Damien Chua, Hong Sheng Cheng, Rachel Tee, Wei Ren Tan, Christopher Ball, Norliza Binte Esmail Sahib, Ser Sue Ng, Jing Qu, Yingzi Liu, Haiyu Hong, Chaonong Cai, Nandini C. L. Rao, Aileen Wee, Mark D. Muthiah, Zoë Bichler, Barbara Mickelson, Jia Qi Lee, Mei Suen Kong, Vanessa S.Y. Tay, Zhuang Yan, Jiapeng Chen, Aik Seng Ng, Yun Sheng Yip, Marcus Ivan Gerard Vos, Debbie Xiu En Lim, Manesh Chittezhath, Jadegoud Yaligar, Sanjay Kumar Verma, Harish Poptani, Xue Li Guan, S. Sendhil Velan, Yusuf Ali, Liang Li, Nguan Soon Tan, Walter Wahli

## Abstract

Adaptive T-cell immune response is essential in conferring protective immunity, a process requiring tight cellular homeostasis regulation. Pathological intrahepatic T-cell landscape has a role in NAFLD propagation; however, its activation remains unknown. To address this gap, we extensively characterized a novel diet-induced NAFLD murine model (LIDPAD) featuring key phenotypic and genetic attributes reflective of human NAFLD. Comparative transcriptomic-guided staging of human and murine NASH reinforced the robustness of LIDPAD in recapitulating critical transitory stages of human NAFLD. We found that angiopoietin-like 4 (Angptl4) shapes activation of the intrahepatic T-cell landscape through the modulation of eIF2α signaling during fibrosis. Single-immune cell analysis and hepatic transcriptomics during fibrosis, and kinase inhibitor screening confirmed that Angptl4 orchestrates the hyperactivation of intrahepatic adaptive immunity via eIF2α signaling. Consistently, immunoblocking of cAngplt4 reduces T-cell overactivation, delaying disease aggravation. Taken together, Angptl4 is a crucial determinant in shaping intrahepatic adaptive immunity during fibrosis in NAFLD.

## INTRODUCTION

Non-alcoholic fatty liver disease (NAFLD) is an impending epidemic that affects close to 30% of the world’s population(*1*). NAFLD encompasses all fatty liver disease states, covering a spectrum of liver conditions associated with metabolic dysregulation, starting with liver steatosis, through nonalcoholic steatohepatitis (NASH) that progresses to cirrhosis(*2*). The global disease prevalence is projected to increase in tandem with the rise of obesity and diabetes mellitus(*3*). Despite the immense burden of NAFLD, there is still no Food and Drug Administration (FDA)-approved pharmacotherapy. The current standard of care is lifestyle modifications targeting weight loss, which is often difficult to achieve and rarely sustainable(*4*). Once the disease has progressed to cirrhosis, such measures become less effective(*5*). Although the challenges and clinical realities have been known for decades, our understanding of the etiology of this multiorgan disease remains sorely lacking, and our ability to treat it remains unacceptably inadequate.

Patients who present with NASH and fibrosis have an increased risk of liver-related mortality(*6*). As liver fibrosis is the strongest predictor of adverse clinical outcomes, NASH represents the best therapeutic condition, during which most clinical symptoms start to manifest. The reversal of liver fibrosis has the greatest clinical impact in the disease and is a key marker in currently defined endpoints for clinical trials. Chronic hepatic inflammation represents the driving force in the progression of NASH to fibrosis/cirrhosis. The evolution of NAFLD in humans and mice is concomitant with an increase in the prevalence of activated cytotoxic CD8^+^ T cells and IFN-γ-producing CD4^+^ T cells in the liver. In addition to T cells, B cells are also detectable within the inflammatory infiltrates in liver biopsies from NASH patients. This hepatic infiltration by B and T cells parallels the worsening of liver damage and lobular inflammation(*7, 8*). Not surprisingly, reduced hepatic recruitment of activated CD4^+^ and CD8^+^ T cells is observed with decrease in fibrosis(*9*). Considering the emerging role of adaptive immunity in the progression of NASH, modulating lymphocyte recruitment and activation offers a novel approach to ameliorate NASH-associated fibrosis. To date, the development of therapies for NASH has mainly focused on modulating metabolic dysfunction, oxidative stress, and innate immunity(*10*). Progress in targeting adaptive immunity in NASH will also rely on a better understanding of the roles of B and T cells in the chronic inflammation process that drives the fibrotic progression in NASH.

Angiopoietin-like 4 (Angptl4) is a secreted glycoprotein belonging to a family of nine structurally similar Angptl proteins(*11*). Initial studies suggested a link between Angptl4 and lipid metabolism(*12–14*). More recently, Angptl4 has been implicated in NAFLD; however, its functions during NAFLD progression remain controversial. Angptl4 upregulates liver cholesterol synthesis via inhibition of lipoprotein lipase (LPL)- and hepatic lipase (HL)-dependent hepatic cholesterol uptake(*15*). Angptl4 is upregulated in liver cirrhosis patients, and whole-body Angptl4 deficiency augments liver fibrosis in a murine methionine-choline deficient diet-induced NASH model through enhanced free cholesterol accumulation in hepatic stellate cells (HSCs)(*16*). On the other hand, *in vivo* overexpression of Angptl4 by an adenoviral vector in high-fat diet (HFD)-fed mice improves glucose tolerance and insulin resistance but induces liver steatosis(*17*). In a mouse cirrhosis model, the suppression of Angptl4 by intravenous injection of sh-Angptl4 lentiviral vector inhibited the polarization and proinflammatory effects of Kupffer cells, the activation of HSCs and fibrosis(*18*). Hepatocyte-specific knockout of Angptl4 protects against HFD-induced glucose intolerance, lipid dysregulation and liver steatosis(*19, 20*). These apparent differences in findings may be attributed to different genetic mouse models and diets. The action of Angptl4 depends on its proteolytic cleavage, releasing an N-terminal coiled-coil domain (nAngptl4) and a C-terminal fibrinogen-like domain (cAngptl4)(*11*). nAngptl4, present in the plasma, is involved in lipid metabolism through inhibitory binding to lipoprotein lipase, curbing peripheral triglyceride release(*21*). cAngptl4 interacts with extracellular matrix proteins and is localized in different tissues(*22*). It has been implicated in various inflammation-associated diseases, such as tissue injury and cancer(*23–25*). cAngptl4 modulates inflammation by enhancing vascular permeability, cytokine secretion, and monocyte differentiation(*26–28*). Whether the domain-specific role of Angptl4 contributes to the abovementioned differences observed in NAFLD, particularly liver inflammation and fibrosis, is unknown, thus currently limiting the development of therapeutics focused on Angptl4 for NASH with fibrosis.

This study aims to improve this situation. First, we describe a physiologically relevant refined diet-induced NAFLD model called LIDPAD that recapitulates key transitory stages of human NAFLD with full-range histological and transcriptomic changes. Second, using the LIDPAD model, we delineated the intrahepatic immune landscape during NASH at single-cell resolution, which revealed the importance of adaptive immunity in disease progression. Finally, we demonstrate that cAngptl4 orchestrates the remodelling of hyperactive adaptive immunity, which aggravates liver inflammation and fibrosis, and show that its inhibition is a promising approach to ameliorate NASH.

## RESULTS

### LIDPAD mice show transitory stages that mirror human NAFLD and extrahepatic complications

Wild-type C57BL/6J male mice housed in the thermoneutral zone (30 °C) were fed either a modified high fat-cholesterol diet of refined ingredients, herein called LIDPAD mice (Liver Disease Progression Aggravation Diet), or a matched Control diet with refined ingredients to ensure comparability (**Table S1**). LIDPAD mice showed a gain in body and liver weight as early as 1 week into the diet, which increased progressively compared with the control group (**Fig. 1A, S1A**). Whole-body calorimetric analysis revealed that LIDPAD mice displayed impaired metabolic flexibility, with less obvious circadian oscillation between carbohydrates and lipid utilization with a stable low respiratory exchange ratio (RER 0.70-0.75), indicating a preference for lipids as fuel compared with control mice (**Fig S2A-B**). The daily energy expenditure shift was also lower in LIDPAD mice when running wheels were introduced, indicating that LIDPAD mice exercise less than control mice (**Fig S2C-E**).

**Figure 1.**
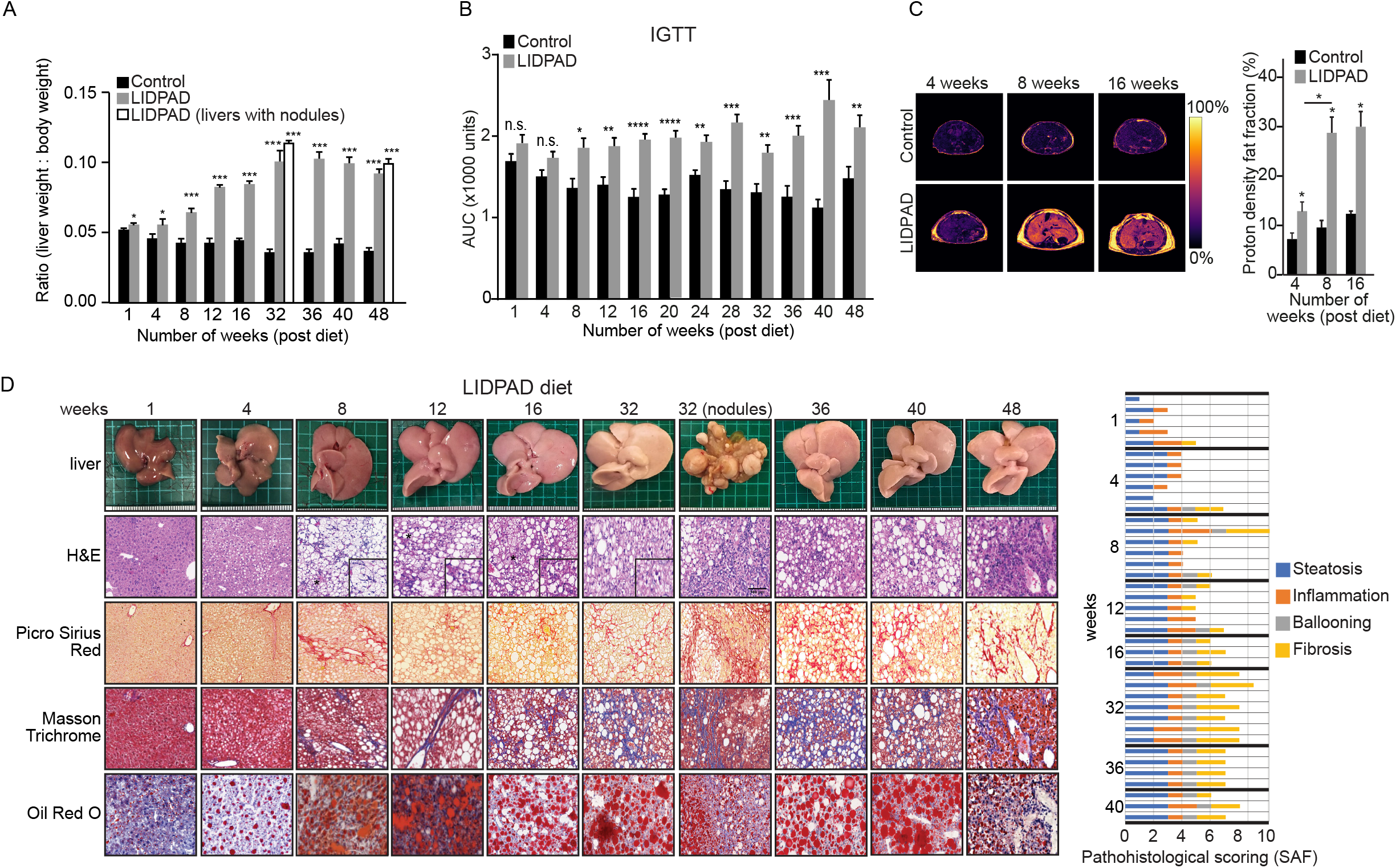
Physiological and metabolic parameters of control and LIDPAD mice. **A.** Liver weight normalized to body weight after feeding for 1 to 48 weeks. **B.** Area under the curve (AUC) of intraperitoneal glucose tolerance test (IGTT) curves for the indicated timepoints. **C.** Proton density fat fraction (PDFF) images from liver of 4-, 8- and 16-week post-diet intervention control and LIDPAD mice. Increased accumulation of hepatic fat is indicated by PDFF scale. Quantification of liver PDFF from control and LIDPAD mice at all imaging timepoints (right). Excessive lipid accretion was observed during 8-12 weeks of LIDPAD intervention. Subcutaneous fat growth identified between timepoints. **D.** Representative macroscopic and microscopic images of the livers obtained from LIDPAD mice. Histological sections were stained with hematoxylin and eosin (H&E) to show general liver features, picrosirius red (PSR), and Masson trichrome (MT) to highlight collagen deposition, and oil red O (ORO) to detect the presence of lipids. The scale bar represents 100 μm. Graphs show tabulated cumulative histological SAF scores of the livers from LIDPAD mice at the indicated weeks postfeeding (right). Each row corresponds to one analysed liver. For 1A-B, n = 7-10 for each group. Data are expressed as the means ± SEMs. ****p<0.0001, ***p<0.001, **p<0.01, *p<0.05 (unpaired t test, ANOVA Welch’s t test or ANCOVA test when appropriate, followed by post hoc comparisons). n.s. denotes not significant. For 1C, n = 5 for each group, *p<0.05 (Mann–Whitney test following pixel thresholding at 60%).

LIDPAD mice developed glucose intolerance within 8 weeks, which corresponded to changes in pancreatic islet size (**Fig 1B, S1B**). The islet area in the control group showed no significant difference from week 1 to 48, with an average islet area of 0.5 μm^2^ ×10^4^ (**Fig S3A-B**). In contrast, in LIDPAD mice, there was a significant increase (1.5-fold) in the islet area between weeks 4 and 16, corresponding to times when glucose intolerance became evident, suggesting compensation via islet cell hyperplasia, consistent with islet compensation during pre-diabetes. From 40 weeks onwards, smaller islets (compared with control mice) started to appear (≤0.5 μm^2^ ×10^4^) in LIDPAD mice, although the islet area remained widely distributed, corresponding to perhaps a deterioration of islet mass because of prolonged insulin resistance, mirroring associations between NAFLD and the development of type-2 diabetes. Nephromegaly characterized by an enlarged interstitial space, increased immune cell infiltration, and mesangial expansion was also observed in LIDPAD mice at 48 weeks (**Fig S3C-D**). Kidney sections also showed fibrosis, suggesting prior renal injury. Given the impaired ability of these mice to metabolize glucose, kidney enlargement could result from diabetic nephropathy, which was not further studied here.

LIDPAD and control mice underwent magnetic resonance imaging-based proton density fat fraction (PDFF) of liver at 4-, 8- and 16-weeks post-diet initiation. At all imaging timepoints, LIDPAD mice had a significantly increased liver fat fraction compared with control mice (**Fig 1C**). LIDPAD-fed mice displayed a hepatic PDFF ranging from 12.9% at 4 weeks post-diet to 30.13% at 16 weeks post-diet. A 2.2-fold increase (12.9 to 28.9%) in the liver fat fraction of the LIDPAD mice was measured between the 4 and 8 weeks imaging timepoints. Hepatic lipid accumulation slowed between 8 and 16 weeks, with the PDFF rising by only 1.3% (**Fig 1C**). Control mice showed an expected slight increase in liver fat fraction across the 4-, 8-, and 16-weeks imaging timepoints, with PDFF ranging from 7.4 to 12.5% (**Fig 1C**).

Histological liver examination remains the gold standard for diagnosing NASH. Liver sections of LIDPAD and control mice at various weeks of diet were scored by pathologists using the Steatosis, Activity, and Fibrosis (SAF) system. Hepatic lipid accumulation was evident as early as 1 week of LIDPAD feeding but was absent in the livers of control mice (**Fig 1D, S4A**), consistent with the MRI imaging data. After 4 weeks, the livers of LIDPAD mice were pale and enlarged with intracellular lipid vacuoles, supported by an SAF score of either grade 2 or 3 in all liver specimens. By 8-12 weeks, mice under LIDPAD developed widespread steatosis, varying levels of intralobular inflammation and infiltration of lymphocytes and macrophages, similar to what is observed in human NASH. In particular, hepatocyte ballooning, the characteristic feature of NASH was seen by 8 weeks. (**Fig 1D, S4B**). Fibrosis was prevalent in 50-80% of the mice (**Fig 1D**). With prolonged feeding of LIDPAD, an increased severity of hepatic fibrosis and more extensive lipid accumulation were observed, except for weeks 32 and 48 (**Fig 1D, S4A**). These instances of reduced steatosis were likely due to decreased lipid accumulation when fibrosis became more prevalent. Liver specimens from weeks 16 through 40 indicated a sustained and consistent increase in all aspects of steatosis, ballooning, inflammation, and fibrosis (**Fig 1D, S4A-B**). From and beyond week 32, hepatocellular nodules were observed in ~13% of the mice, comparable to the rate of ~2-12% for human hepatocellular carcinoma(*29*).

Altogether, LIDPAD mice display a wide array of physiological and physical activity impairments. The LIDPAD mouse model presents the entire spectrum of NAFLD disease progression, with steatosis starting in the early timepoints (weeks 1-4) and NASH appearing after 8 weeks. Fibrosis was noted in 80% of mice starting at 12 weeks. A proportion of LIDPAD mice eventually develop endstage sequelae such as cirrhosis and hepatocellular nodules, mirroring NAFLD patients in disease progression and incidence.

### Hepatic transcriptomic analyses and cytokine profiling highlight hepatic inflammation and fibrosis

To identify key cellular mechanisms that underpin the observed histological changes in NAFLD, we performed transcriptomic and biochemical analyses. Hierarchical clustering of differentially expressed genes (DEGs) across all LIDPAD liver samples revealed three gene groups according to their expression trends with diverse Gene Ontology profiles (**Fig 2A-B**). Among the prominent early events were the dysregulation of cholesterol and lipid homeostasis, as well as the acute-phase response, which indicates hepatic inflammation in response to lipotoxicity. LIDPAD mice also have a hepatic transcriptomic profile of altered cellular response to LPS between weeks 1-4, suggesting a compromised gut barrier. As NASH developed, genes involved in collagen assembly and inflammatory response pathways appeared and intensified, concurring with the histology and pathological scoring for NASH (**Fig 2A-B**).

**Figure 2.**
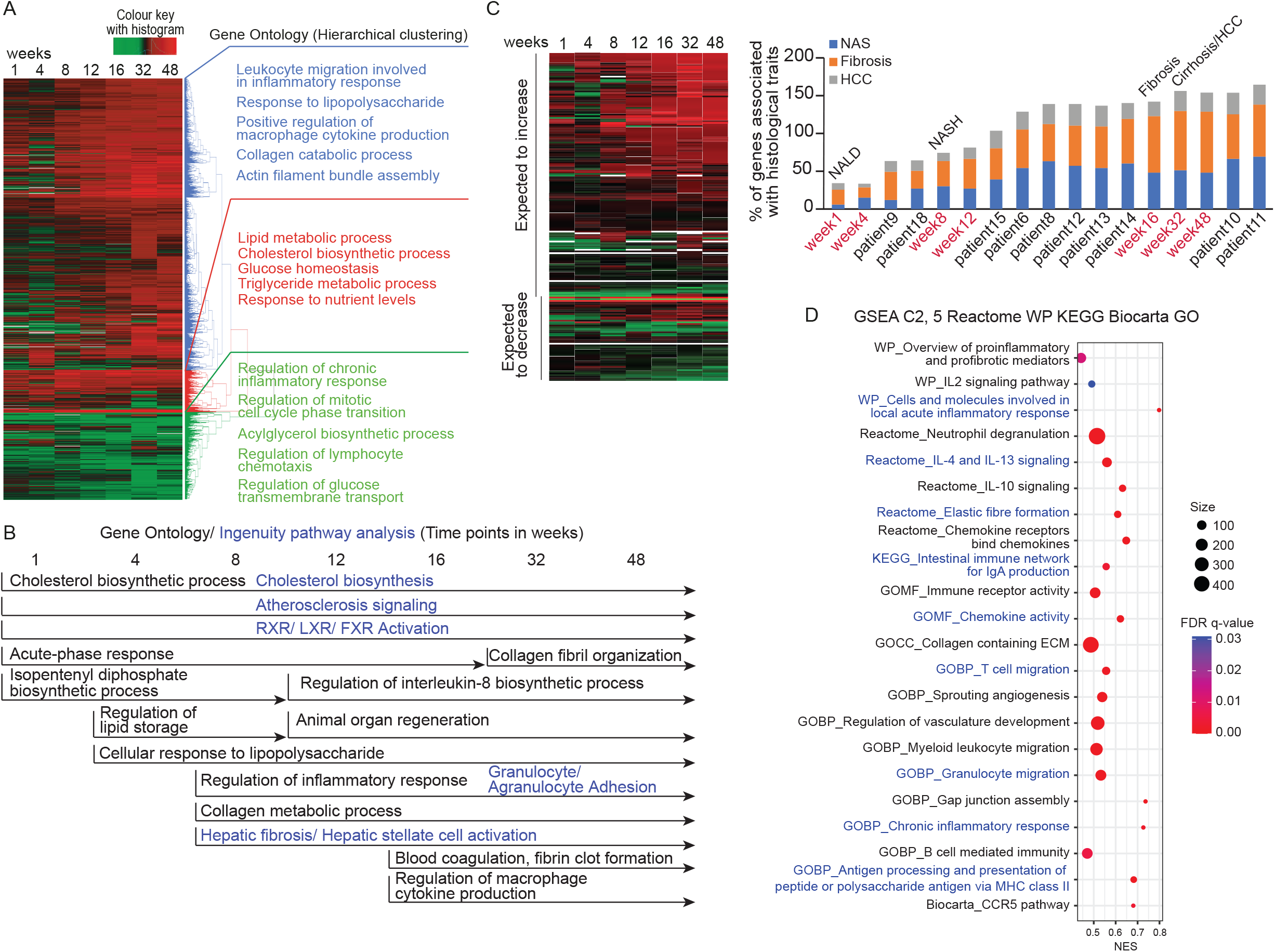
Hepatic transcriptomic and molecular analysis of control and LIDPAD mice. **A.** Heatmap of differentially expressed genes (DEGs) across weeks 1 to 48 with corresponding Gene Ontology (GO) biological process terms guided by hierarchical clustering. **B.** Gene ontology and pathway analysis across multiple time points tracking temporal changes in DEGs, featuring only recurring GO terms that span across at least two time points, to increase the confidence of the data as a true positive. **C.** Transcriptomic profile of LIDPAD mice plotted with orthologues of human genes associated with NAFLD progression. Percentage of genes associated with each histological trait for patients with NASH. Graphs are arranged with increasing severity (left to right). Transcriptomic-based staging of LIDPAD livers was included as a reference. **D.** GSEA plotted using FDR and size, showing terms from Reactome, WP, KEGG, Biocarta and GO.

We confirmed our above findings with a series of biochemical measurements. We observed a significant increase in alanine transaminase (ALT) levels, as a biomarker of liver damage, in LIDPAD mice beginning at week 8 of the diet (**Fig S5A**). Although there was a mild increasing trend of ALT in control mice during aging, this increase was not statistically significant. Corroborating the transcriptomics analysis, both free cholesterol and squalene levels were elevated in the LIDPAD mice compared to the controls, as determined by LC–MS (**Fig S5B**). A basal amount of cholesterol was detected in the livers of control mice, a site for cholesterol synthesis, whereas squalene levels were close to none (**Fig S5B**). The increase in free cholesterol can be attributed to cholesterol in the LIDPAD diet. The elevation in the concentration of squalene, a precursor of cholesterol, suggests that the sterol biosynthetic pathway was affected by LIDPAD.

Various studies have identified serum cytokines in the evaluation of NAFLD patients(*30*). Multiplex immunoanalysis of serum revealed that no single cytokine remained elevated throughout the progression of steatosis to cirrhosis. We observed distinct yet overlapping early-, mid-, and late-stages of cytokine profiles (**Fig S5C**). At weeks 1 to 4, we observed an increase in low-grade chronic inflammatory cytokines (IL-6 and IL-13) and inflammatory chemokines (MIP-1α and eotaxin), consistent with the start of steatosis and liver damage. At weeks 8-12, the inflammatory response was sustained by a new group of cytokines (TNF-α and IL-1α) and chemokines (MCP-1 and MIG) that further promoted the infiltration of immune cells (see below). After 12 weeks, the early- and mid-stage cytokines decreased and were replaced by another group of cytokines that are important for the maturation of adaptive immune cells (IL-7, IL-9, IL-10, and IL-2) (**Fig S5C**). While cohort studies have identified similar serum biomarkers, they failed to address the temporal changes as disease progresses. Thus, different panels of cytokines/chemokines appear useful for monitoring the progression of NAFLD.

Next, we compared our LIDPAD model results to a human meta-analysis of NAFLD transcriptomic signatures of 218 genes associated with histological worsening(*31*). The expression of most genes detected in the LIDPAD model exhibited the same-sign consistency compared to human gene expression (**Fig 2C**). Guided by histological examination and major shifts in temporal gene expression, we revealed that the weeks 1, 8, 16, and 32 of LIDPAD feeding corresponded to key transition phases of human NAFLD progression (**Fig 2C**). We performed transcriptomic-guided staging of 10 NASH patients to assess the robustness of the LIDPAD transcriptomic progression signature, which revealed that most of the patient’s liver biopsies (6 out of 10) corresponded to weeks 12 to 16 of our LIDPAD mouse model, i.e., NASH with fibrosis (**Fig 2C**). Histological evaluation of matched liver specimens showed some degree of concordance based on ordinal SAF staging (**Fig S5D**). Gene set enrichment analysis (GSEA) of our human NASH transcriptomes revealed that biological processes such as hepatic inflammation, fibrosis and vascular perturbations predominated in human NASH livers (**Fig 2D**).

### Angptl4 deficiency alters the intrahepatic immune cell landscape to delay liver fibrosis

To understand the role of Angptl4 in the immunopathogenesis of fibrosis (see Introduction), we examined the effect of Angptl4 deficiency in two models: myeloid cell-specific Angptl4 mutant mice (Angptl4^LysM−/−^) and mice treated with a neutralizing antibody (mAb) against cAngptl4 during LIDPAD-induced NASH(*28, 32*). Mice at 4 weeks of LIDPAD feeding were treated with mAb. These mice continued to gain weight similar to control mice. mAb did not alter glucose intolerance induced by LIDPAD feeding (**Fig S6A-B**). Similarly, Angptl4^LysM−/−^ mice gained weight but had significantly higher glucose tolerance across all conditions. These observations suggest that immunoneutralization of cAngptl4 does not affect glucose disposal following bolus administration, whereas genetic Angptl4 deletion in myeloid cells improves whole body glucose tolerance.

At 8 weeks of LIDPAD, wild-type mice develop steatosis and inflammation in the liver, with increased prevalence and severity of NASH and fibrosis by week 12. Notably, both the mAb treatment and Angptl4^LysM-/-^ mice had reduced hepatic inflammation and fibrosis (**Fig 3A-B, S6C-E**). Hepatic transcriptomic profiles confirmed the ameliorative effects of targeting Angptl4 to delay NASH progression. Principal component analysis (PCA) revealed that the liver transcriptomes of mAb-treated LIDPAD mice occupy the space between control and LIDPAD mice, with diminished differences between mAb-treated LIDPAD mice and LIDPAD mice on longer feeding, suggesting a slower disease onset in the former (**Fig 3C**). The hepatic transcriptomes of Angptl4^LysM-/-^ mice formed a distinct cluster from other groups that shifted toward LIDPAD mice with prolonged feeding, suggesting a different hepatic impact from the combined effects of gene ablation and high-calorie feeding. DEG analysis identified five functional clusters (**Fig 3D**). As expected, the LIDPAD diet markedly activated the hepatic inflammatory response and ECM remodelling activities, which became stronger with prolonged feeding. These proinflammatory and profibrotic events were effectively diminished by cAngptl4 mAb, leading to slower progression. While most of the NASH-related immune response was suppressed in Angptl4^LysM-/-^ mice, different sets of proinflammatory genes that were associated with fibrinolysis, complement activation, and the adaptive immune system were triggered in these mice, highlighting the interrelationship between innate and adaptive immune responses. Our analysis revealed an immunomodulatory role of cAngptl4 and the benefits of targeting cAngptl4 to delay diet-induced NASH.

**Figure 3.**
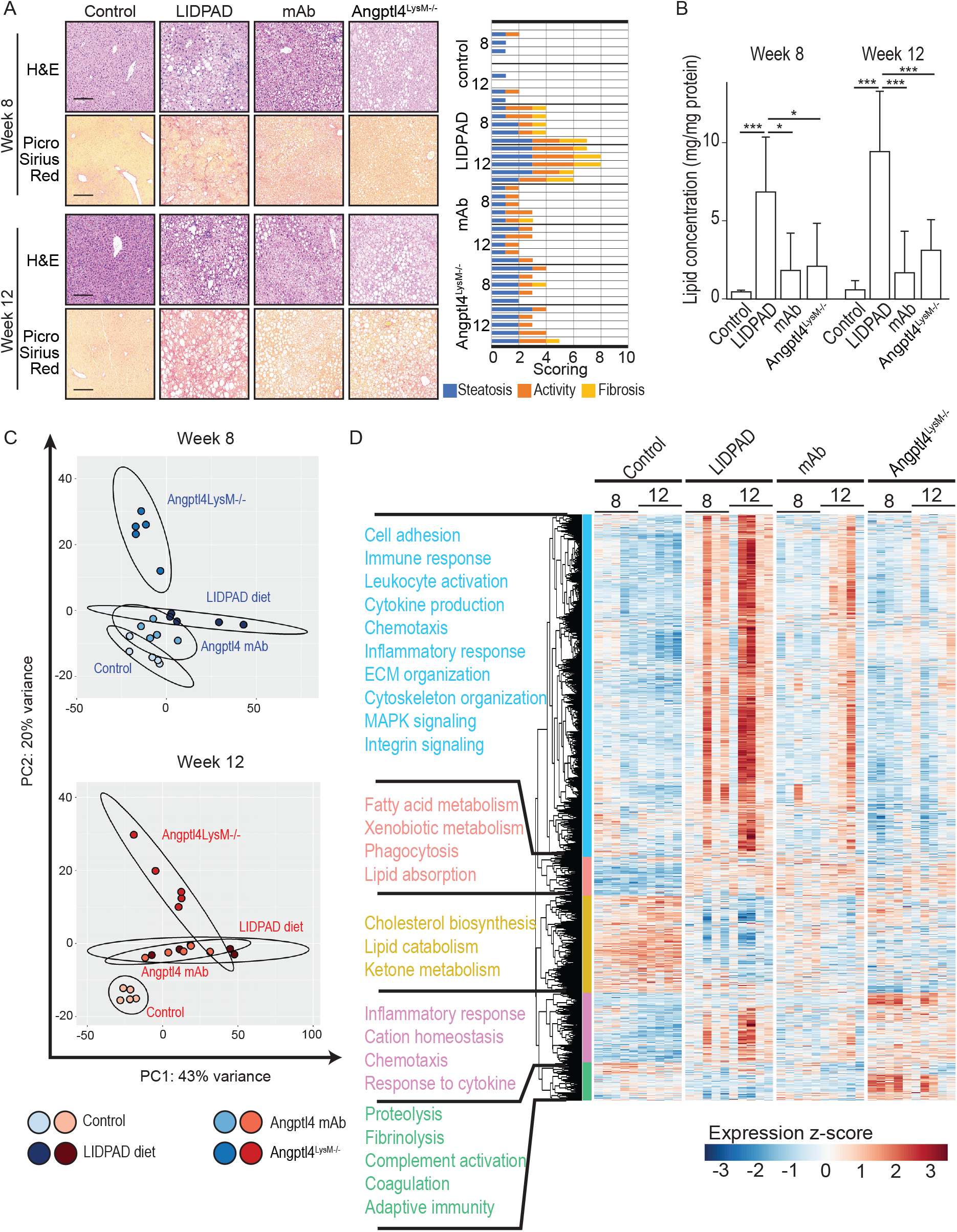
Effects of Angptl4 deficiency on liver pathology and transcriptomes. **A.** Representative images of microscopic images of the livers obtained from control, LIDPAD-fed mice, LIDPAD-fed mice treated with neutralizing mAb against cAngptl4 and LIDPAD-fed Angptl4^LysM-/-^ mice after 8 and 12 weeks of feeding. Histological sections were stained with hematoxylin and eosin (H&E) to show general liver features and picrosirius red (PSR) to highlight collagen deposition. The scale bar represents 100 μm. The graphs show the tabulated cumulative histological SAF scores of the livers from the indicated groups at the indicated weeks postfeeding (right). Each row corresponds to one analysed liver. **B.** Graph showing the hepatic triglyceride concentration normalized to the protein level. Data are expressed as the means ± SEMs. ***p<0.001, *p<0.05 (unpaired t test). n = 5 for each group. **C.** Principal component analysis (PCA) of liver transcriptomes across the indicated groups and treatments at 8 and 12 weeks. **D.** Heatmap of differentially expressed genes (DEGs) at weeks 8 and 12 with corresponding Gene Ontology (GO) biological process terms guided by hierarchical clustering of hepatic transcriptomes of the indicated groups and treatments.

### Single-cell transcriptomics reveals adaptive immune cell remodelling during fibrosis

Immuno-deficient NSG mice developed steatosis with only mild fibrosis after chronic (1 year) LIPDAD feeding, confirming the essential prominent role of immune cells in fibrosis (**Fig S6F**). To gain insight into the immune cell landscape as NASH livers develop fibrosis, scRNA-seq of intrahepatic immune cells from LIDPAD and control livers was performed at weeks 8 and 12. Nineteen immune cell subpopulations were identified, with B and T lymphocytes constituting most of the intrahepatic immune cell population. B lymphocytes (differentiated and matured plasma cells) and T lymphocytes (8 major cell subclusters) made up ~43.2% and ~41.2% of the total intrahepatic immune cells, respectively. Six subpopulations of myeloid-derived immune cells, namely, neutrophils, monocytes, Kupffer cells, monocyte-derived macrophages (mo-macs), classical dendritic cells (cDCs) and plasmacytoid DCs (pDCs), accounted for close to 10% of the CD45^+^ cells in the livers (**Fig 4A-B, Table S2**).

**Figure 4.**
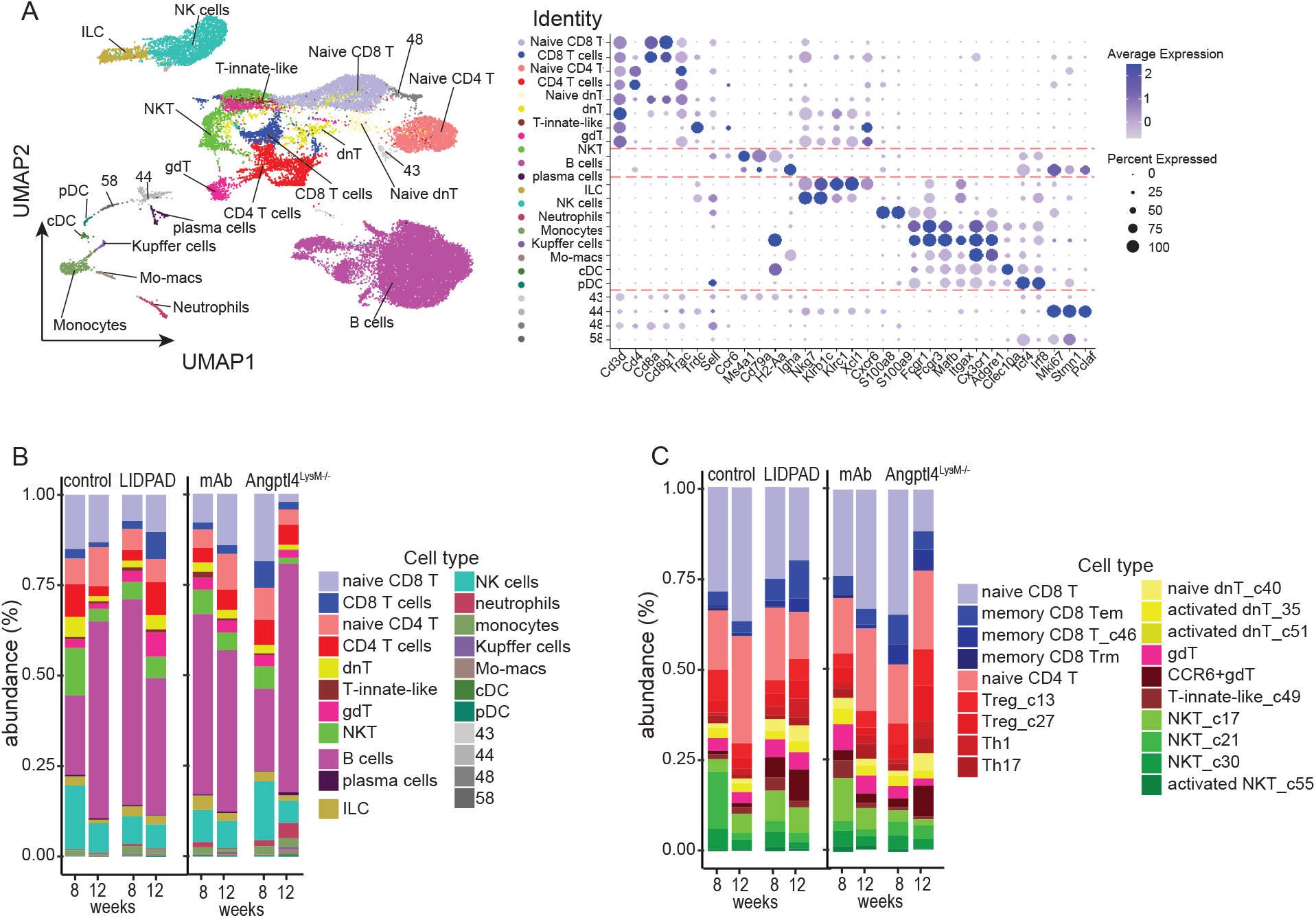
Intrahepatic immune landscape during LIDPAD-induced NAFLD. **A.** Overall UMAP representation of the total intrahepatic CD45^+^ immune population (left). Dotplot representing molecular markers used to identify major immune cell types (right). **B-C.** Relative abundance of total CD45^+^ immune cell types (B) and T-cell populations (C) in the livers of control mice, LIDPAD-fed mice, LIDPAD-fed mice treated with neutralizing monoclonal antibodies against cAngptl4 (mAb) and LIDPAD-fed Angptl4^LysM-/-^ mice after 8 and 12 weeks of feeding.

Most myeloid-lineage cells were marginally increased in NASH livers at weeks 8 and 12 compared with the control diet (**Fig 4B, Table S2**). On the other hand, B lymphocytes increased at week 8 and decreased at week 12. The proportion of CD8^+^ T, CD4^+^ T, dnT, and NKT cells showed an opposite trend compared with the control diet, i.e., decreased at week 8 and increased at week 12, suggesting their involvement in fibrosis. High-resolution analysis of the T-cell subpopulations revealed 4 subclusters of CD8^+^ T cells, 5 subclusters of CD4^+^ T cells, 3 subclusters of dnT cells, 5 subclusters of innate-like T and NKT cells, and 2 clusters of gdT cells (**Fig 4C, S7A, Table S3**). CD4^+^ and CD8^+^ T cells demonstrated dynamic shifts in proportions across conditions, accounting for approximately two-thirds of the T-lymphocyte population (**Fig 4C**). We interrogated the fate of T-cells to further understand their role. As the disease progressed from 8 to 12 weeks, more naïve CD4^+^ T cells were activated, as evidenced by scRNA-sequencing and FACS analysis (**Fig 5A-C, S8**). Pseudotime analysis revealed two major peaks along the CD4^+^ T-cell differentiation trajectory that correspond to the activation state (**Fig 5B**). Similar shifts in intrahepatic CD8^+^ T cells were also observed (**Fig S9A-C**). In summary, fibrosis progression is marked by remodelling of the lymphoid-lineage immune cell landscape, particularly an increase in activated CD4^+^ and CD8^+^ T cells.

**Figure 5.**
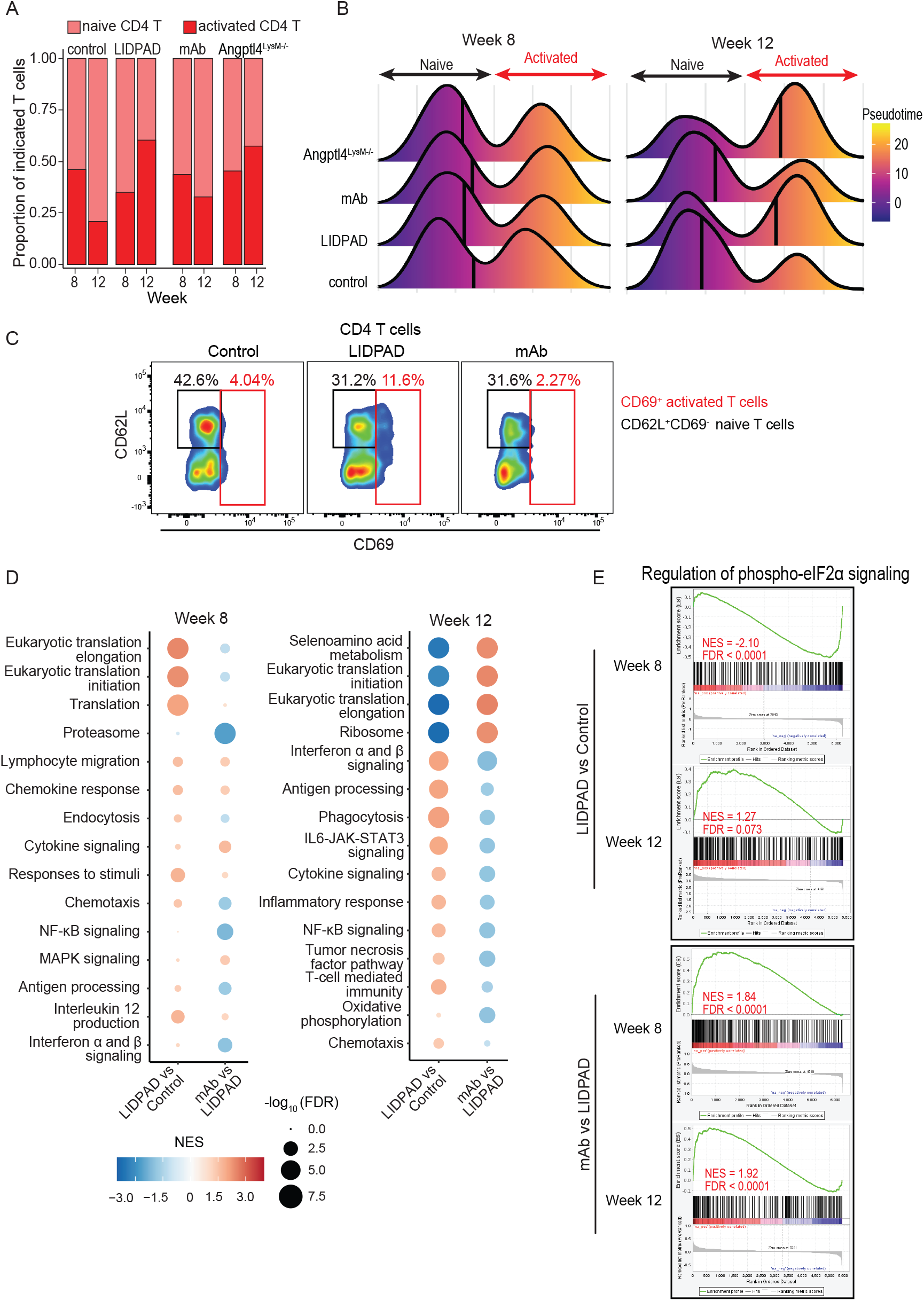
cAngptl4 mAb suppressed intrahepatic CD4 T-cell activation via eIF2α-mediated translation attenuation. **A.** Proportion of intrahepatic naïve and activated CD4 T cells in control mice, LIDPAD-fed mice, LIDPAD-fed mice treated with neutralizing monoclonal antibodies against cAngptl4 (mAb), and LIDPAD-fed Angptl4^LysM-/-^ mice after 8 and 12 weeks of feeding. **B.** Pseudotime analysis ordering the CD4 T cells according to the activation status. Vertical lines indicate the median pseudotime score of each group. **C.** Percentage of intrahepatic naïve (CD45^+^Lin^-^CD3+CD4+CD62L+CD69^-^) and activated (CD45^+^Lin^-^CD3^+^CD4^+^CD69^+^) CD4 T cells from the indicated groups after 12 weeks of feeding based on FACS. Lineage markers (Lin) include CD11b, NK1.1 and Ly6G. **D-E.** GSEA showing highly enriched gene sets in CD4^+^ T cells during NASH progression from 8 to 12 weeks of feeding **(D)** and signature genes involved in the regulation of eIF2α phosphorylation **(E)** based on the differential transcriptomes of naïve CD4 T cells from different treatment groups. Normalized enrichment score (NES) > 0 and < 0 indicate the enrichment of the gene set by up- and downregulated genes, respectively. For 5D, the NES is color-coded, and the dot size denotes the −log10 FDR of representative enrichment tests.

In LIDPAD-fed Angptl4^LysM-/-^ mice, there was a higher neutrophil persistence and macrophage:monocyte ratio than in LIDPAD-fed mice, suggesting a prolonged innate inflammatory response. Our study also revealed an interrelationship between innate and adaptive immune responses during fibrosis. The directionality of change in the relative abundance of T and B cells was opposite to LIDPAD mice and similar to control mice, i.e., total B cells increased, and T cells decreased from week 8 to 12. (**Table S2**). However, no change in the activation status of CD4^+^ and CD8^+^ T cells was observed compared with LIDPAD-fed mice. The mAb treatment of LIDPAD-fed mice reduced the abundance of many immune cells, e.g., neutrophils, mo-macs and monocytes, to levels similar to the control mice. mAb treatment attenuated the relative abundance of T and B cells from weeks 8 to 12 compared with LIDPAD mice (**Table S2–S3**). Importantly, and in contrast to Angptl4^LysM-/-^ and LIDPAD-fed mice, mAb treatment effectively blocked the increase in activated CD4^+^ and CD8^+^ T cells from weeks 8 to 12 (**Fig 5A-C, S9A-C**). Hence, the activation of CD4^+^ and CD8^+^ T cells is a key event in fibrosis, which was diminished by blocking cAngptl4.

GSEA of CD4^+^ and CD8^+^ T cells revealed gene sets in two primary processes, namely, eukaryotic translation and immune responses (**Fig 5D, S9D**). During fibrosis, we observed a downregulation of eukaryotic translation-associated genes, accompanied by a concomitant upregulation of genes involved in the inflammatory response in CD4^+^ T cells (**Fig 5D)**. In CD8^+^ T cells, the eukaryotic translation gene sets became insignificant by week 12, while the genes involved in cell recognition and cytokine-mediated responses were upregulated (**Fig S9D**). These observations suggest that the regulation of eukaryotic translation activities is pivotal for the proinflammatory phenotypes of the two T-cell lineages. mAb treatment repressed eukaryotic translation machineries in CD4^+^ and CD8^+^ T cells, accompanied by substantial inhibition of inflammatory activities in both CD4^+^ and CD8^+^ T cells at week 12 (**Fig 5D, Fig S9D-E**). The effect of mAb on eIF2α signaling was observed in CD8^+^ T cells at week 8 (**Fig S9E**), which is likely due to the complex activation mechanism of CD8^+^ T cells that require inputs from CD4^+^T helper cells. Using a curated gene set from StringDB and Reactome, we identified a stimulatory effect of the cAngptl4 mAb on the phosphorylated eukaryotic initiation factor-2α (phospho-eIF2α)-mediated signaling, which is often associated to global translation attenuation and could contribute to the suppressed T-cell activation in mAb-treated mice (**Fig 5E**).

### cAngptl4-mediated eIF2α signaling modulates the activation of CD4^+^ T cells

Phosphorylation of eIF2α reduces global translation, allowing cells to conserve resources and to initiate a reconfiguration of gene expression to effectively manage stress conditions. We sought to decipher how cAngptl4 modifies phospho-eIF2α signaling. Active phospho-eIF2α signaling stimulates the expression of three canonical target genes, activating transcription factor 4 (ATF4), heat shock protein family A member 5 (HSPA5), and DNA damage inducible transcript 3 (DDIT3). As protein kinases are ubiquitously involved in cellular signaling activities, we conducted kinase inhibitor screens to map the regulatory pathways of Angptl4 on eIF2α phosphorylation (based on the readouts from target gene expression) using LPS-activated wild-type and Angptl4^-/-^ CD4^+^ T cells. Unsupervised hierarchical clustering of the results revealed 3 clusters of kinase targets corresponding to (i) uninvolved kinases, (ii) Angptl4-independent kinases, and (iii) Angptl4-dependent kinases (**Fig 6A**). In wild-type cells, a quarter of the kinase inhibitor has minimal effect on the expression of these target genes compared with DMSO treatment, indicating that these kinases are not involved in phospho-eIF2α-mediated signaling. In DMSO-treated Angptl4^-/-^ cells, target gene expression was already higher than in DMSO-treated wild-type cells, suggesting that Angptl4 deficiency activates phospho-eIF2α-mediated signaling of these target genes. As expected, inhibition of these uninvolved kinases did not alter the target genes compared to DMSO-treated Angptl4^-/-^ cells (**Fig 6A-B**).

**Figure 6.**
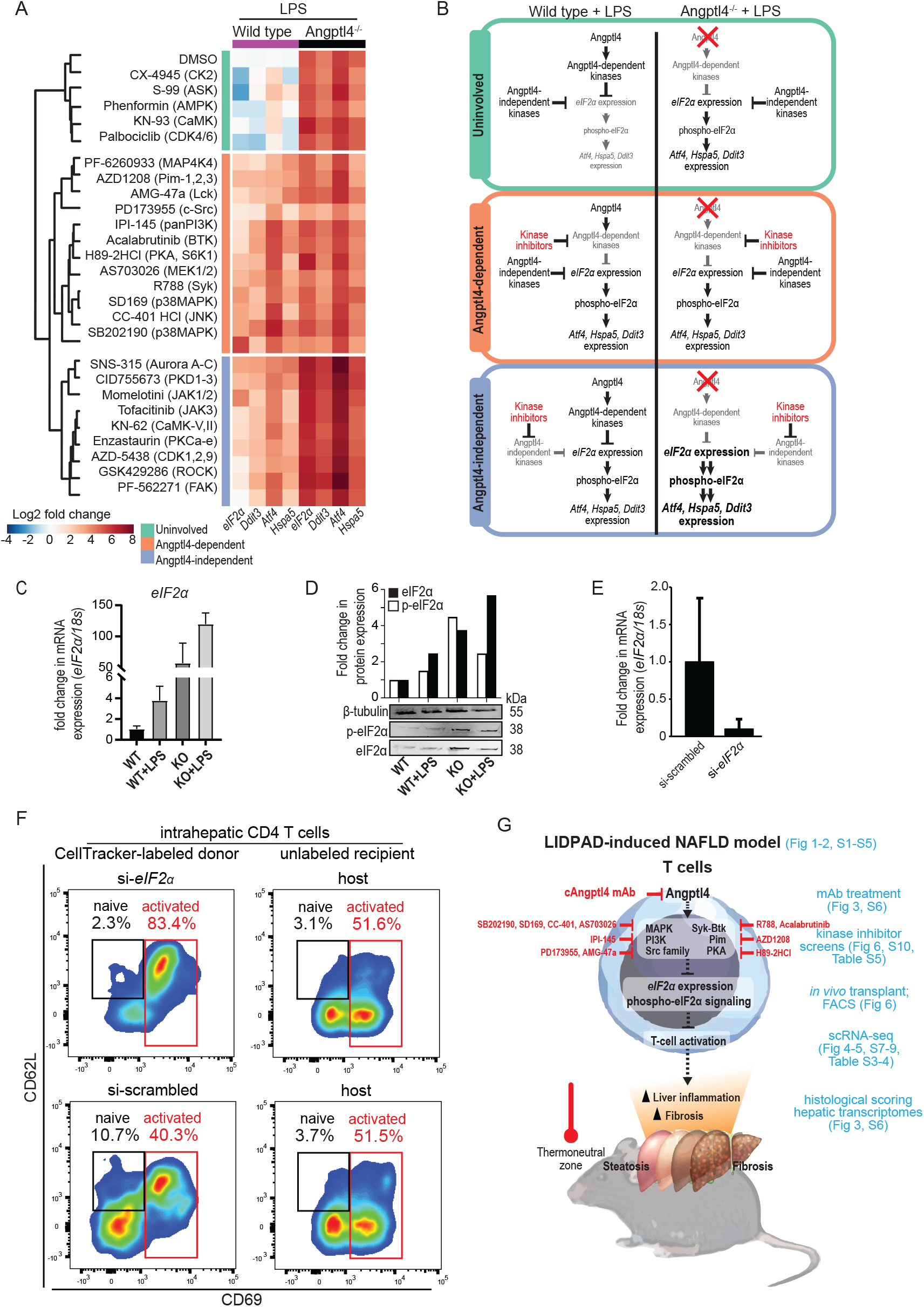
Kinomic modulatory pathway of the Angptl4-eIF2α phosphorylation pathway. **A-B.** Kinase inhibitor screens of CD4^+^ T cells from wild-type and Angptl4-KO mice using LPS as a stimulant. The heatmap illustrates the expression levels of *eIF2α* and target genes of phosphorylated-*eIF2α* (i.e., *Ddit3, Atf4* and *Hspa5*) relative to DMSO-treated wild-type CD4^+^ T cells **(A)**. The kinase inhibitors and their molecular targets (in parenthesis) were separated into three groups with differential activities on phospho-eIF2α signaling and Angptl4 dependency based on the expression profiles using a hierarchical clustering method. The schematic diagram **(B)** outlines six scenarios to justify the role of different protein kinases in the Angptl4-eIF2α axis. Terms and arrows that are shrunken and greyed out indicate suppression of their activities, while those that are enlarged indicate overactivity. Briefly, Angptl4 deficiency in CD4^+^ T cells is associated with upregulation of *eIF2α* and phospho-*eIF2α* target genes, suggesting an inhibitory effect of Angptl4 on *eIF2α* expression and its downstream signaling. For protein kinases that do not regulate the *eIF2α* pathway, their inhibitors do not modify the expression of *eIF2α* and the target genes in wild-type and Angptl4-knockout CD4^+^ T cells (*e.g*., CK2, ASK, AMPK, CaMK and CDK4/6). When an inhibitor upregulates *eIF2α* and the target genes in wild-type but not Angptl4-knockout CD4+ T cells (which reflects derepression of *eIF2α* in the absence of Angptl4), the regulatory activity of the corresponding molecular targets on the eIF2α pathway is Angptl4-dependent (*e.g*., MAPK, PI3K, Src, Syk-Btk, etc.). For kinase inhibitors that further upregulate *eIF2α* and the target genes regardless of the Angptl4 status, the regulatory activity of molecular targets is Angptl4-independent (*e.g*., PKD1/2/3, JAK1/2/3, Aurora A/B/C, etc.). **C-D.** mRNA expression of *eIF2α* **(C)** and immunoblots of eIF2α and phospho-eIF2α **(D)** in wild-type and Angptl4-knockout CD4^+^ T cells with and without LPS challenge. For **(C)**, *18s* was used as the endogenous reference gene while for **(D)**, β-Tubulin which served as a loading control, was from the same samples. **E-F.** Knockdown efficiency of *eIF2α* in Angptl4-knockout mouse splenic lymphocytes using SMARTpool siRNA **(E)**. *18s* was used as the endogenous reference gene. Activation status of Cell Tracker-labelled, adoptively transferred (top) and host (bottom) CD4^+^ T cells in the livers of LIDPAD-fed mice after 12 weeks of feeding **(F)**. Adoptively transferred cells were treated with either si-*scrambled* or si-*eIF2α* at 10 nM. Naïve (CD45^+^Lin^-^CD3+CD4+CD62 L+CD69^-^) and activated (CD45^+^Lin^-^CD3^+^CD4^+^CD69^+^) CD4 T cells were determined using FACS. **G.** Schematic diagram summarizing the Angptl4-eIF2α pathway and its role in T-lymphocyte activation during fibrosis in LIDPAD-induced NAFLD mouse model. Angptl4 stimulates the kinases that inhibit phospho-eIF2α signaling, removing its inhibitory effect on T cell activation, i.e., a reduced phospho-eIF2α signaling is associated with more T cell activation.

Twenty-one kinase inhibitors activated the expression of target genes in wild-type cells, indicating that these kinases repress phospho-eIF2α-mediated signaling (**Fig 6A-B**). Among them, nine kinase inhibitors against Angptl4-independent kinases derepressed phospho-eIF2α-mediated signaling, increasing the expression of the target genes. These inhibitors further elevated the expression of target genes in Angptl4^-/-^ cells compared with DMSO treatment. The remaining 12 kinase inhibitors did not alter the expression of the target genes in Angptl4^-/-^ cells compared with DMSO-treated Angptl4^-/-^ cells and were identified as Angptl4-dependent (**Fig 6A-B**). Angptl4 deficiency did not activate Angptl4-dependent kinases, and futile inhibition by the respective kinase inhibitors yielded minimal changes in the expression of target genes (**Fig 6A-B**). Generally, the expression profile of eIF2α followed those of the target genes, revealing a positive correlation between eIF2α and phospho-eIF2α signaling. Angptl4^-/-^ cells had higher eIF2α expression than wild-type cells, which was further increased by LPS **(Fig 6C)**. Immunoblot analyses showed higher phosphorylated eIF2α in Angptl4^-/-^ CD4 T cells than in wild-type cells, regardless of LPS exposure **(Fig 6D)**. The results confirmed a suppressive effect of Angptl4 on phospho-eIF2α through the inhibition of eIF2α expression and our kinase inhibitor screens reveal three predominant kinase pathways, i.e., MAPK, PI3K, and Src, in Angptl4-eIF2α axis.

To confirm the *in vivo* role of phospho-eIF2α signaling in CD4^+^ T-cell activation, we monitored the effect of eIF2α deficiency on the fate of Angptl4^-/-^ CD4^+^ T cells, which has high endogenous eIF2α, in LIDPAD mice. CellTracker-labelled donor Angptl4^-/-^ CD4^+^ T cells treated with si-eIF2α were introduced intravenously into LIDPAD recipient mice (**Fig 6E**). As expected, an increase in activated host CD4^+^ T cells (unlabelled) was detected in the livers of LIDPAD-fed mice. The deficiency of eIF2α in labelled donor CD4^+^ T cells significantly increased the percentage of activated CD4^+^ T cells compared to si-scrambled donor CD4^+^ T cells (**Fig 6F**). Evidently, a reduced *eIF2α* expression suppresses phospho-eIF2α signaling, leading to more T cell activation. Taken together, we systematically showed the pivotal role of Angptl4-eIF2α signaling in the activation of T cells that contributes to increased inflammation and fibrosis (**Fig 6G)**.

## DISCUSSION

The pathogenesis of NAFLD is a multifactorial and multistep process that involves ill-characterized gene-environment interactions and multiorgan involvement(*33*). In this study, we performed single-cell analysis of the intrahepatic immune landscape of a human relevant diet-induced NAFLD mouse model to reveal an indispensable role of adaptive immune cell remodelling in disease aggravation characterized by increased activated T-lymphocytes. We showed that the activation of CD4+ T cells depends on eIF2α-mediated eukaryotic translation initiation during fibrosis. We further decipher an Angplt4-eIF2α signaling pathway essential for the activation of CD4^+^ T cells (**Fig 6G**). The immunoneutralization of cAngptl4 effectively diminishes the NASH-promoting immunological response and delays liver fibrosis.

Understanding the network of multisystem interactions requires a physiologically relevant animal model that can mirror the many human NASH features and comorbidities(*34*). An improved diet-inducible liver disease mouse model was established using a refined Liver Disease Progression Aggravation Diet (LIDPAD). Species differences between the LIDPAD model and humans were minimized via thermoneutral mouse housing conditions, while refined diets mitigate batch-to-batch compositional variations, which are metabolic confounders. One key feature of the NAFLD model is its fast disease progression that can capture the full range of anthropometric, physiologic, histological, and transcriptomic anomalies of human pathogenesis in a considerably shorter time frame, i.e., simple steatosis in 1-4 weeks, chronic inflammation in 4-8 weeks, and fibrosis in 12-16 weeks. LIDPAD mice rapidly gain weight and develop impaired glucose homeostasis. Noninvasive longitudinal imaging through magnetic resonance techniques and temporal hepatic transcriptomic profiling pointed to the shift from fatty liver to NASH at week 4 of the diet and the development of fibrosis at 12-16 weeks. Using transcriptomic-guided staging of patients with biopsy-confirmed NASH emphasizes the relevance of LIDPAD mice to model human NAFLD. This model supports a liver-pancreas axis that promotes compensatory pancreatic islet cell hyperplasia, a phenomenon similarly observed in animal models of insulin resistance(*35*) and in humans, where there is a clear correlation between BMI and β-cell mass(*36*) and thereafter fatty liver and the development of type-2 diabetes(*37*). Furthermore, LIDPAD mice also displayed NAFLD-associated vasculopathy, with heightened endothelial expression of CXCL12 in the aortas and the liver vasculature(*38*). These various metabolic dysfunctions and extrahepatic comorbidities are collectively absent in current NAFLD models.

Understanding the time frame for the fatty liver to NASH transition in the LIDPAD model allows for in-depth examination of the role of immune cells in disease aggravation to fibrosis. Histological examination revealed elevated infiltration of cells from the adaptive immune system at 8 weeks of LIDPAD feeding, which persisted in a more advanced stage. Our single-cell surveillance of intrahepatic immune cells of livers in LIDPAD mice at this transition revealed an overrepresentation of adaptive immune cells, including B- and T-lymphocytes. These abnormal immune responses are predominantly orchestrated by CD4^+^ helper T cells, which are highly activated throughout the early and advanced NASH stages in the LIDPAD model. This corroborates findings that T-lymphocytes are heavily involved in NAFLD deterioration(*38–40*). A recent study also identified an integral role of activated CD4^+^ T cells at fibrotic sites in NASH-driven hepatic inflammation and fibrosis(*41*). Mechanistically, our study revealed that Angptl4-eIF2α signaling is pivotal for the activation of CD4^+^ T cells. CD4^+^ T cells also mediate NASH exacerbation by modulating the activation of CD8^+^ T cells, NKT cells and fibrogenic HSCs. Mechanistically, CD8^+^ cytotoxic T cells drive the NAFL-NASH transition through elevated proinflammatory cytokine secretion and autoaggressive killing of hepatocytes(*9, 42*). Different CD4^+^ T-cell subsets exhibit diverse effects on liver fibrosis. Th2- and Th17-derived cytokines stimulate collagen production and TGFβR-dependent hepatic fibrosis(*43*). Notably, NAFLD patients also exhibit increased serum proinflammatory and profibrogenic cytokine signatures coupled with increased activated CD4^+^ T cells(*43*). Collectively, the multifaceted role of the CD4^+^ T-cell response remains an integral driver in NASH exacerbation.

Previous studies emphasized the lipid metabolism function of Angptl4 in NASH and yielded contradictory conclusions. In addition to the dynamic changes in immune cell landscapes during disease progression, the different functions of nAngptl4 and cAngptl4 may have obscured our full systems understanding of the protein. Our study particularly focused on the exacerbation of NASH with fibrosis and affirmed an immunomodulatory role for cAngptl4 in the intrahepatic adaptive immune response through the Angptl4-eIF2α axis. While active phospho-eIF2α signaling limits ER stress, hepatocyte death and fibrogenesis in diet-induced NASH(*44, 45*), very little is known about its role in the immunopathology of NAFLD. We showed that active eIF2α supports the maturation and clonal expansion of CD4^+^ T cells, which subsequently mediate the activation of other lymphocytes to create an inflamed and fibrogenic liver environment in NASH. Indeed, inhibition of the components in the translation initiation complex abrogates CD4^+^ T-cell activation and differentiation(46), which was also observed in our NAFLD mice bearing eIF2α-knockdown CD4^+^ T cells via adoptive transfer. Our study also showed reduced fibrosis in myeloid-specific Angptl4-knockout mice, likely due to reduced T-cell infiltration rather than their activation status. Angptl4 has been shown to affect the mobilization efficiency of hematopoietic stem/progenitor cells from bone marrow and the maturation of specific myeloid cell populations(*47–49*). The immunomodulation of T cells offers an exciting avenue for alleviating chronic inflammation that contributes to NASH. For example, the oral administration of anti-CD3 (foralumab) promotes the induction of Treg cells, alleviates insulin resistance, suppresses the chronic inflammation associated with NASH and exerts a beneficial effect on clinically relevant parameters(*50*). We show that immunotherapy with administration of mAb against cAngptl4 effectively diminishes T-lymphocyte activation and delays liver fibrosis in NASH via modulation of phosphorylated-eIF2α signaling.

In summary, we described a human relevant diet-induced NAFLD model, thus offering opportunities to reliably translate findings in mice to insights into human NAFLD for drug development. Using this model, we confirmed a domain-specific role for cAngptl4 and phospho-eIF2α signaling in curbing the activation of T-lymphocytes, especially CD4^+^ T cells, during fibrosis.

## MATERIALS AND METHODS

### Animals, feeding regimens and treatment

Male wild-type C57BL/6J (Invivos, Singapore), LysMCre-*Angptl4^-/-^* and NOD scid gamma (NSG, Jackson Laboratory) mice aged between 8-9 weeks were given *ad libitum* access to LIDPAD (Liver Disease Progression Aggravation Diet, modified from Teklad diet TD. 88137, Envigo, USA) or a control diet (Teklad Custom Diet, modified from AIN-93M, Envigo, USA) (**Table S1**) and water. LysMCre-*Angptl4^-/-^* mice have site-specific knockout of *Angptl4* expression, primarily in myeloid immune cells. Neutralizing monoclonal antibodies against mouse cAngptl4 (clone 3F4F5) were administered intraperitoneally at 10 mg/kg twice weekly after 4 weeks of NAFLD induction with LIDPAD. Animals were kept in standard housing cages for up to 48 weeks at a controlled temperature and humidity of 30 °C (thermoneutral zone) and 49.9%, respectively, with a 12-hour darklight cycle. Body weight and food consumption were measured weekly. Animals were euthanized using CO2 prior to blood and tissue collection. Organs were fixed with 4% PFA or snap frozen with liquid nitrogen. All experiments were carried out following the guidelines of the Institutional Animal Care and Use Committee (IACUC) (SingHealth IACUC: #2014/SHS/1008; NTU-IACUC: A18031, A10032, A18033, A18042, A20055).

### Patient samples

Ten NAFLD patients and 5 liver cancer patients scheduled for liver surgery in the Fifth Affiliated Hospital of Sun Yat-Sen University were recruited to provide surplus liver tissue excised from the surgery. The patients had no prior or existing liver infection or other active infectious diseases. NAFLD liver tissue samples were obtained from the NAFLD regions of the 10 NAFLD patient livers. Normal liver tissue samples were obtained from the normal tissue surrounding the liver tumors of the 5 liver cancer patients. The obtained samples were then cut in halves and kept either in formaldehyde for histological analysis or in a −80 °C freezer before RNA extraction. The study was approved by the Institutional Review Board of the Fifth Affiliated Hospital of Sun Yat-Sen University, Zhuhai, China (Approval No. L136-1), and written consent was obtained from all the participants.

### Liver histopathology

Liver samples were fixed in 4% paraformaldehyde and embedded in paraffin. Histological sections were cut into 5 μm thick sections, dewaxed, and rehydrated before staining with H&E, PSR, and MT via a Leica Autostainer XL (Leica, Germany). Frozen tissue samples for ORO staining were prepared and stained following an isopropanol-based protocol. Images were captured using Axioscan. Z1 (Zeiss, Germany) under brightfield settings at 20x magnification. A blinded histological assessment was performed using the Steatosis, Activity, and Fibrosis (SAF) scoring system described in the Supplementary Methods.

### RNA sequencing

Total RNA was extracted from frozen liver samples obtained from all mouse groups and patient samples with TRIzol reagent (Life Technologies, U.S.) and E.Z.N.A.^®^ HP Total RNA kit (Omega Biotek, U.S.). RNA samples were sequenced using the Illumina HiSeq platform to generate 50-bp paired-end reads. Raw reads were mapped to the *Mus musculus* genome assembly from Ensembl, GRCm38 or GRCh38 for human(*51*), via HISAT2(*52*). Uniquely mapped reads were analysed with FeatureCounts(*53*) to produce gene count matrices, which were subjected to differential expression analysis using DESeq2(*54*). Genes meeting the criteria of >±1 log2-fold-change and adjusted p value <0.05 were considered differentially expressed genes (DEGs). DEGs were stratified by expression and analysed using Gene Ontology and Ingenuity Pathway Analysis (QIAGEN Inc.) to identify the top-ranked enriched pathways. Raw sequences were deposited into GEO with accession numbers GSE159911 and GSE214504.

### Single-cell isolation of intrahepatic immune cells

Intrahepatic immune cells were isolated as previously described with some modifications(*55*). Briefly, mice were anesthetized with 100 mg/kg ketamine and 10 mg/kg xylazine, injected with 3 μg of anti-mouse CD45-FITC antibodies (Miltenyi Biotec, Germany) retro-orbitally and left for 3 min prior to euthanasia by cardiac puncture. The livers were harvested and dissociated into single-cell suspensions using the Liver Dissociation Kit on a GentleMACS Octo Dissociator (Miltenyi Biotec). The cell suspension was filtered through a 70 μm strainer. Lymphocytes in the supernatant were enriched with Percoll density gradient centrifugation (1.07 g/mL) and treated with RBC lysis buffer to deplete red blood cells. The lymphocyte fractions were stained with anti-CD45-APC (panhematopoietic cell marker) and propidium iodide (Live/Dead stain) before sorting on a FACSAria Fusion cell sorter (BD Biosciences, USA). Live intrahepatic immune cells (CD45-FITC-CD45-APC+ PI-) were collected, washed twice, and counted before single-cell library preparation. Single-cell RNA sequencing and annotation of the scRNAseq immune subpopulation are described in the Supplementary Methods.

### FACS

Intrahepatic immune cells were isolated as mentioned above. Single cells were obtained through a 70 μM strainer, blocked with 3% BSA containing FcR blocker and stained with antibodies (**Table S4**). The proportions of naïve (CD45^+^Lin^-^CD3^+^CD4^+^CD8^-^CD62L^+^CD69^-^) and activated (CD45^+^Lin^-^CD3^+^CD4^+^CD8^-^CD69^+^) CD4^+^ T cells, and naïve (CD45^+^Lin^-^CD3^+^CD4^-^CD8^+^CD62L^+^CD69^-^) and activated (CD45^+^Lin^-^CD3^+^CD4^-^CD8^+^CD69^+^) CD8^+^ T cells were analysed with LSRFortessa X-20 (BD Biosciences, USA) based on the gating strategy in **Fig. S8**.

### Kinase inhibitor screen

Splenic CD4^+^ T cells were isolated from wild-type and Angptl4^-/-^ murine lymphocytes (methodology as above) using a CD4^+^ T-cell magnetic separation column (Miltenyi Biotech). Isolated CD4^+^ T cells were cultured in AIM-V culture media (Thermo Fisher Scientific, USA) supplemented with 5% wild-type and Angplt4^-/-^ mouse serum. CD4^+^ T cells were treated with a panel of 28 kinase inhibitors (SYN-2103; SYNkinase, Victoria, Australia & TargetMol, Massachusetts, USA) at the stipulated concentrations (**Table S5**) overnight before treatment with 1 μg/mL LPS for 2 h. The log2(fold change) relative expressions of eIF2α, and target genes of eIF2α-mediated downstream genes (i.e., DDIT3, HSPA5 and ATF4) of each kinase inhibitor treatment were compared against LPS-treated wild-type conditions via qPCR analysis.

### Adoptive transfer

Splenic lymphocytes were harvested from Angptl4-knockout mice as described above. Knockdown of eIF2α (or *Eif2s1*) was performed using 10 nM of Accell mouse *Eif2s1* SMARTpool siRNA (Horizon Discovery, UK) based on our published protocol(*28*). Control cells were transfected with scrambled siRNA. After knockdown, the lymphocytes were labelled with CellTracker^™^ Green CMFDA Dye (Thermo Fisher Scientific, USA) at 1 μM for 45 mins. A total of 1×10^7^ cells were intravenously injected into mice fed the LIDPAD diet for 12 weeks to create a NASH environment. Intrahepatic immune cells were isolated to examine the proportion of naïve and activated cells in donor (labelled with Cell Tracker Green; FITC^Hi^) and recipient (unlabelled; FITC^Lo^) CD4^+^ T cells via FACS.

## ACKNOWLEDGEMENT

This research is supported by the following:

NTUitive SPARK program (WW), Lee Kong Chian School of Medicine, Nanyang Technological University Singapore Start-Up Grant (WW), Ferring Singapore Innovation grant (WW, NS), Singapore Ministry of Education under its Singapore Ministry of Education Academic Research Fund Tier 1 (RG30/20) (NS), Grant 81900071, National Natural Science Foundation of China (LL), Singapore Ministry of Education under its Singapore Ministry of Education Academic Research Fund Tier 2 (MOE2017-T2-1-038) (YA). Intramural funding support from the Institute of Bioengineering and Bioimaging, Agency for Science Technology and Research (V.S.S.)

Z.S.L. and D.C. are recipients of the LKCMedicine PhD scholarship and Nanyang Presidential Graduate scholarship, respectively, under N.S.T. supervision. A.S.N. is recipient of the National Science Scholarship from the Agency for Science, Technology, and Research. C.H.S. is a recipient of LKCMedicine Dean’s Fellowship. J.Q.L. is supported by the CN Yang Scholars Programme of NTU Singapore. X.L.G. is a recipient of the 2016 Nanyang Assistant Professorship from NTU. C.B. is an ARAP programme awardee.

We wish to acknowledge Dr. Hervé Guillou [Toxalim (Research Center in Food Toxicology), INRAE] for his critical reading and suggestions. We also wish to thank the support of LKCMedicine for the creation and execution of the NAFLD Investigation Centre (N.I.C.E.).

## AUTHOR CONTRIBUTIONS

W.W. conceived the Liver Disease Progression Aggravation Diet (LIDPAD) project; S.S.N., B.M., W.W. developed the LIDPAD; S.S.N. conducted the proof-of-concept experiments; R.T. organized the animal experimentation; Z.B. and W.W. conceived and designed the metabolic and behavioral analysis; D.C., C.H.S. and N.S.T. conceived, designed, and performed the T-cell and Angplt4 experiments. D.C., C. H.S., M.I.G.V., Y.S. and N.S.T. conceived, performed, and analysed the single-cell RNA-sequencing experiments. C.B., J.Y., S.K. V, H.P. and S.S.V conducted MRI imaging and PDFF analysis. Z.S.L., D. C., C.H.S., R.T., W.R.T., N.B.E.S., J.Q.L., M.S.K., V.S.Y.T., Z.Y., J.C., A.S.N., Y.S.Y., M.I.G.V., D.X.E.L and M.C. performed the animal experiments; Z.S.L, W.R.T., D.C. and H.S.C. performed all sequencing data analyses. A.W., N.C.L., M.D.M. scored the liver biopsies as clinical pathologists. M.S.K. prepared and analysed the histology of the kidney. V.S.Y.T. and Y.A. prepared and analysed the histology of the islet cells. X.L.G. processed and analysed the lipids. J. Q., Y. L., H. H., C.C., and L.L. performed histological staining, scoring and RNA sequencing on the patient samples. Z.S.L., D.C., C.H.S., N.S.T and W.W. wrote the manuscript with inputs from all authors.

## DATA AND MATERIALS AVAILABILITY

High-throughput sequencing data can be found in the GEO database (GSE159911, GSE214504 and GSE214172). All other data needed to evaluate the conclusions in the paper are present in the paper and/or the Supplementary Materials.

---

**Table S1.**
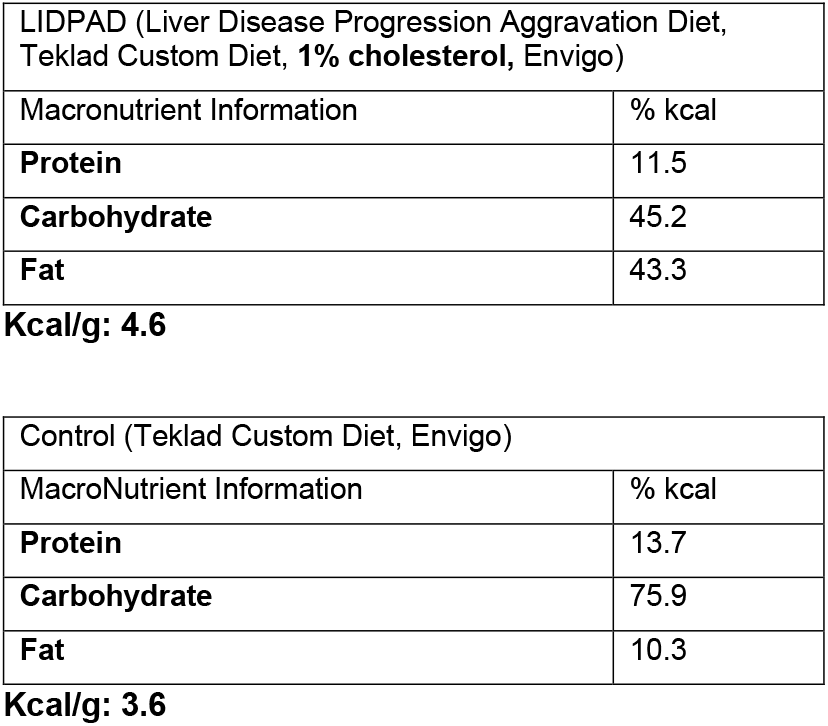
Nutritional Information on Diets (Experiment)

**Table S2.**
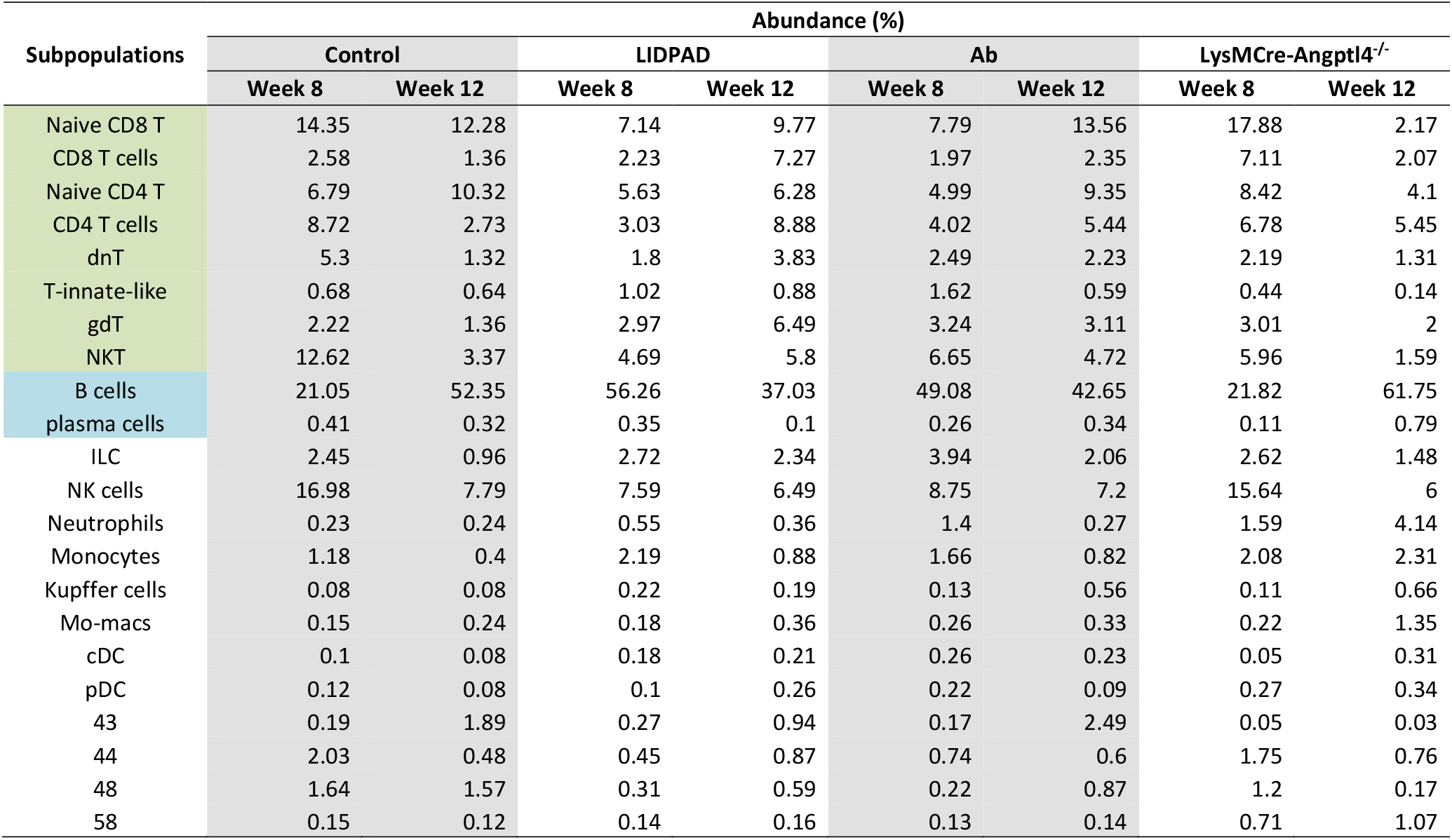
Relative abundance of intrahepatic immune cell subpopulations.

**Table S3.**
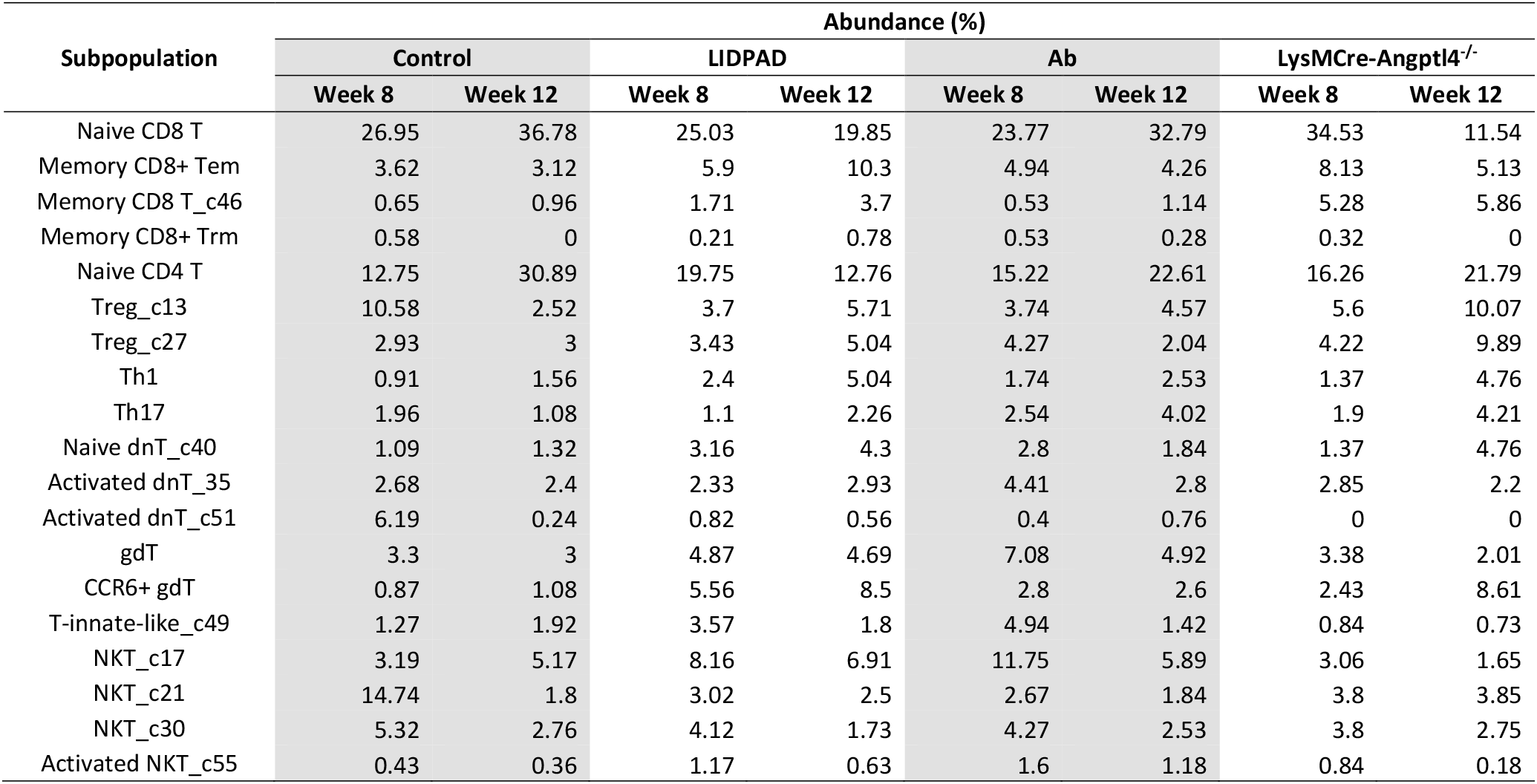
Relative abundance of T cell subpopulations.

**Table S4.**
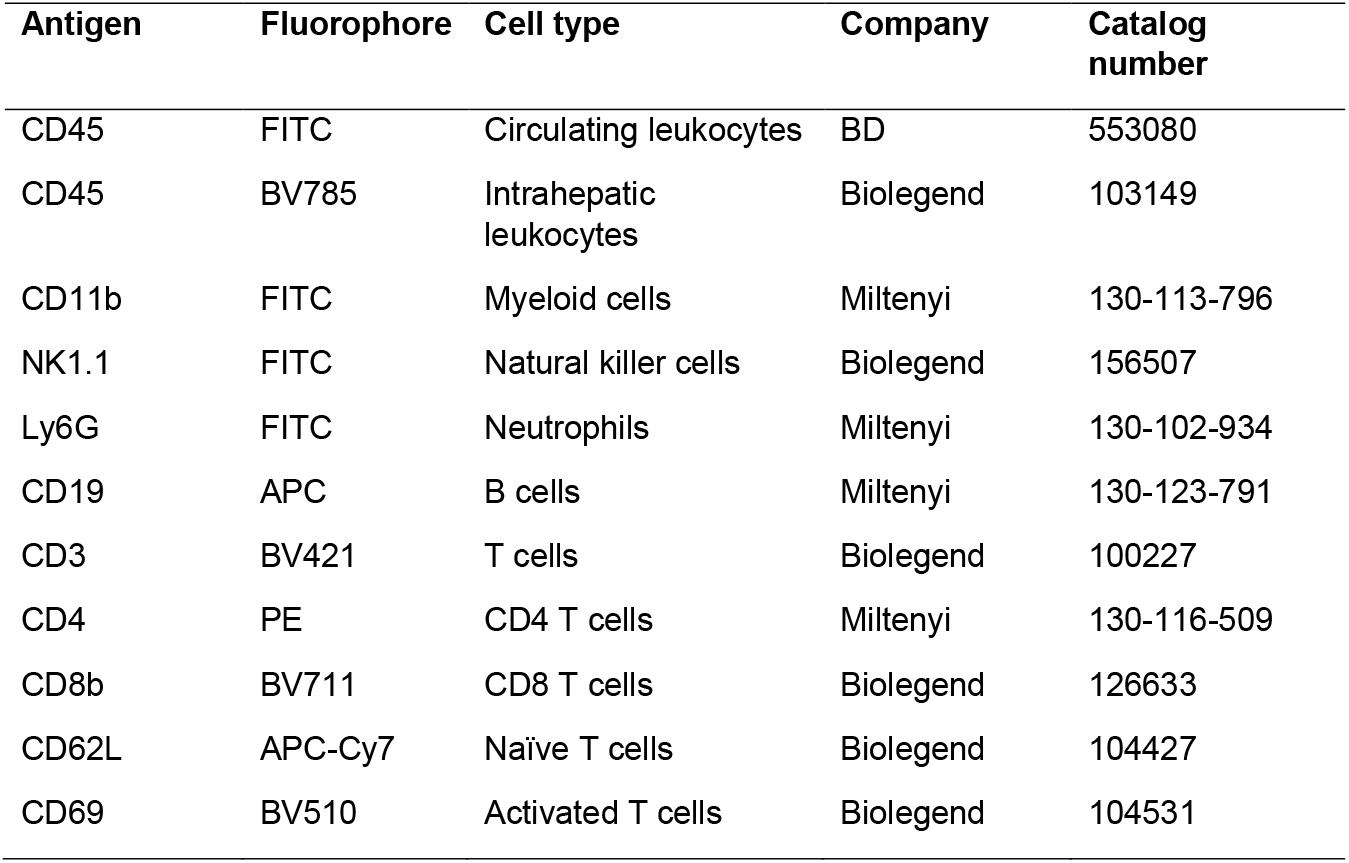
Antibodies used for FACS.

**Table S5.**
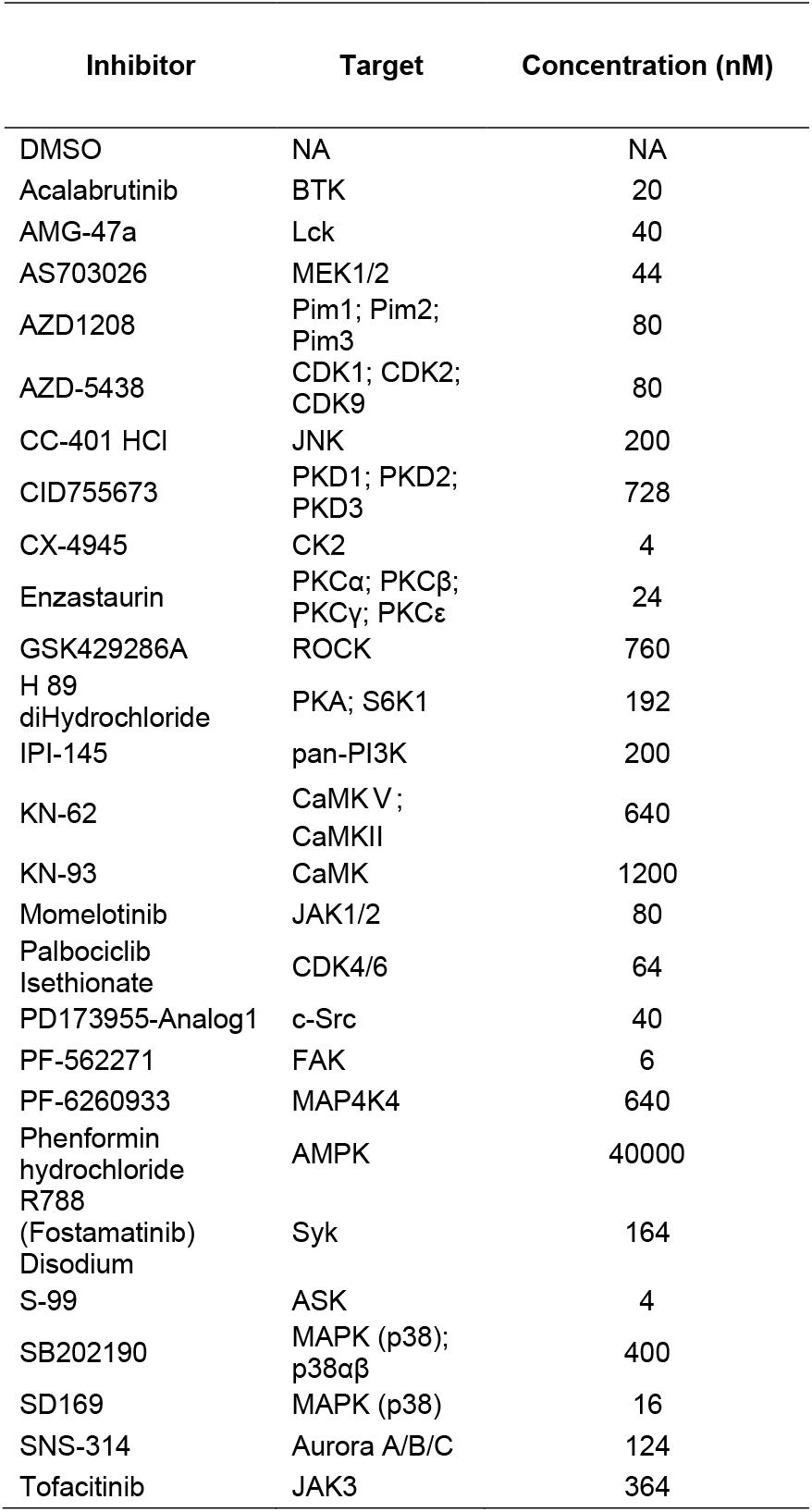
List of kinase inhibitors used, molecular targets and concentration used in the drug screen.

## Supplemental Figure Legends

**Figure S1:**
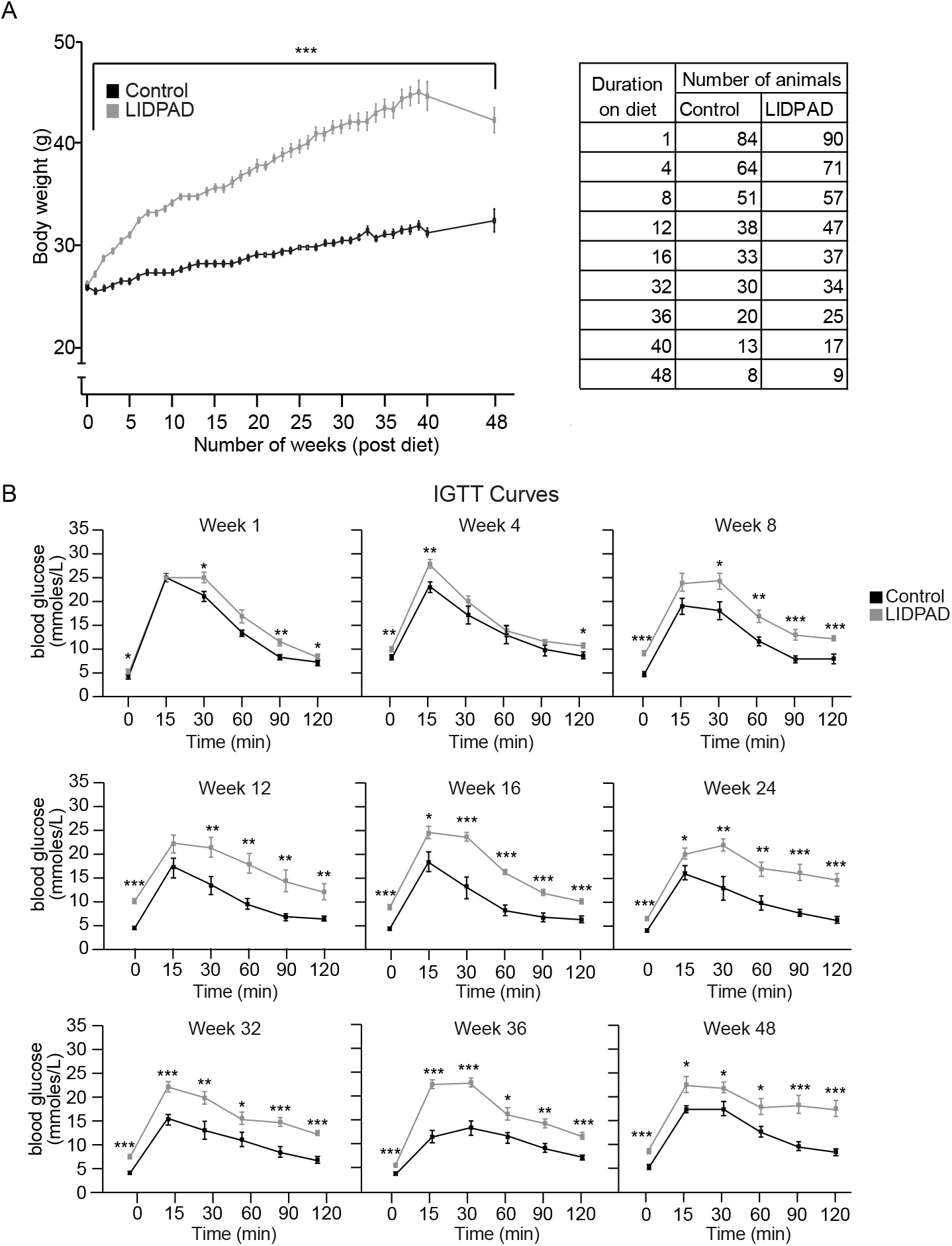
Metabolic parameters of control and LIDPAD mice. **A.** Change in weight (left) of mice fed the control diet and LIDPAD and the number of animals (left) measured at each time point for 48 weeks. **B.** IGTT curves of LIDPAD- and control-fed mice from weeks 1 to 48. n = 7-10 per group. Data are expressed as the means ± SEMs. ****p<0.0001, ***p<0.001, **p<0.01, *p<0.05 (unpaired t test, ANOVA Welch’s t test or ANCOVA test when appropriate, followed by post hoc comparisons).

**Figure S2:**
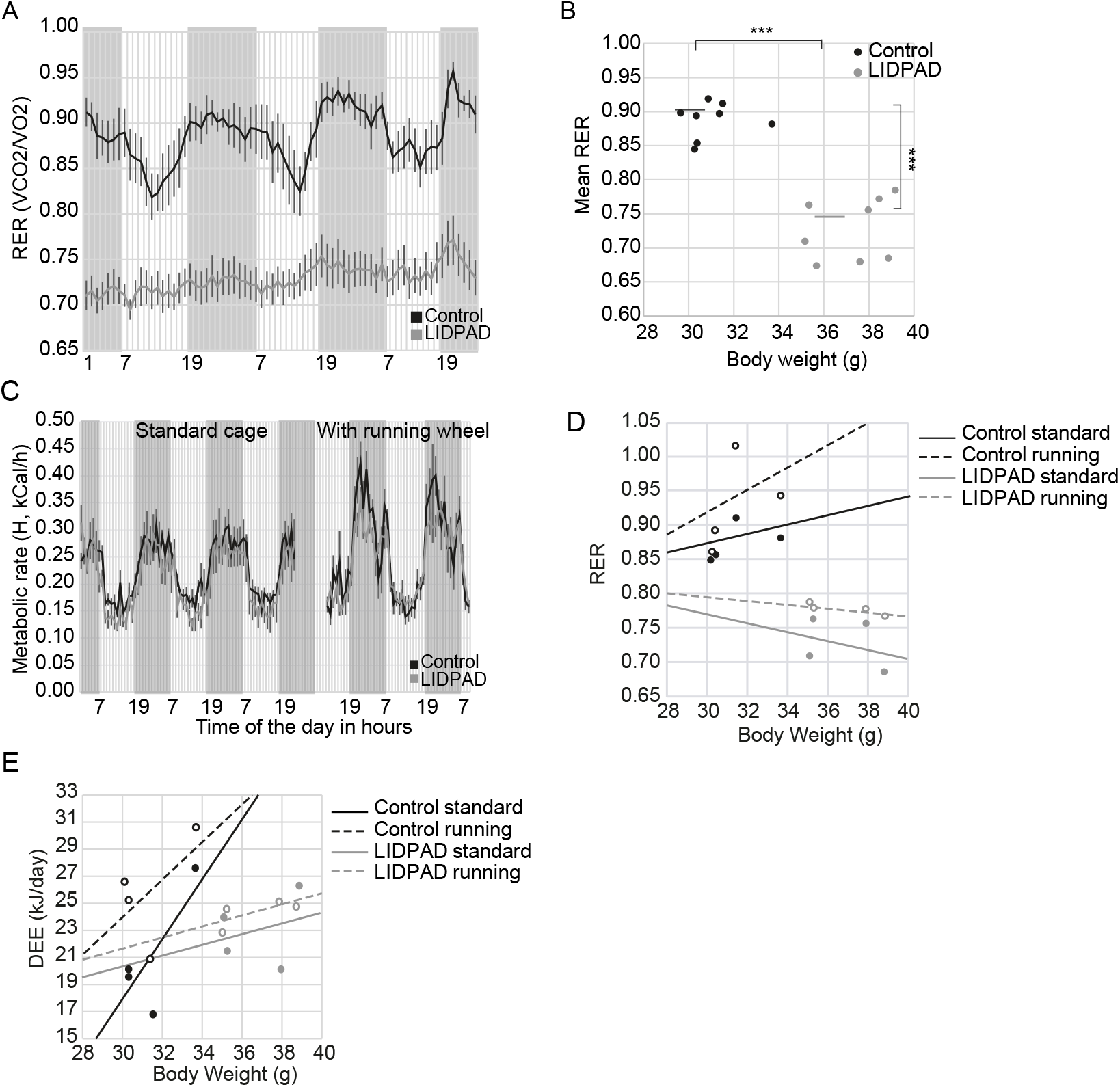
Physiological parameters of control and LIDPAD mice. **A-B.** Respiratory exchange ratio (RER) **(A)** of mice fed the control and LIDPAD diets every hour. Gray and white zones represent dark and light cycles that alternate from 0700 to 1900 daily. Mean RER **(B)** was calculated to determine any statistical differences between control and LIDPAD mice. **C-E.** Mouse physiology was recorded when challenged with voluntary exercise. Metabolic rate **(C)** displayed with gray and white zones representing dark and light cycles. The respiratory exchange ratio **(D)** of LIDPAD mice remained below 0.8, and the daily energy expenditure **(E)** of LIDPAD mice remained relatively unchanged after introducing voluntary exercise. Data are expressed as the means ± SEMs. ***p<0.001 (unpaired t test).

**Figure S3:**
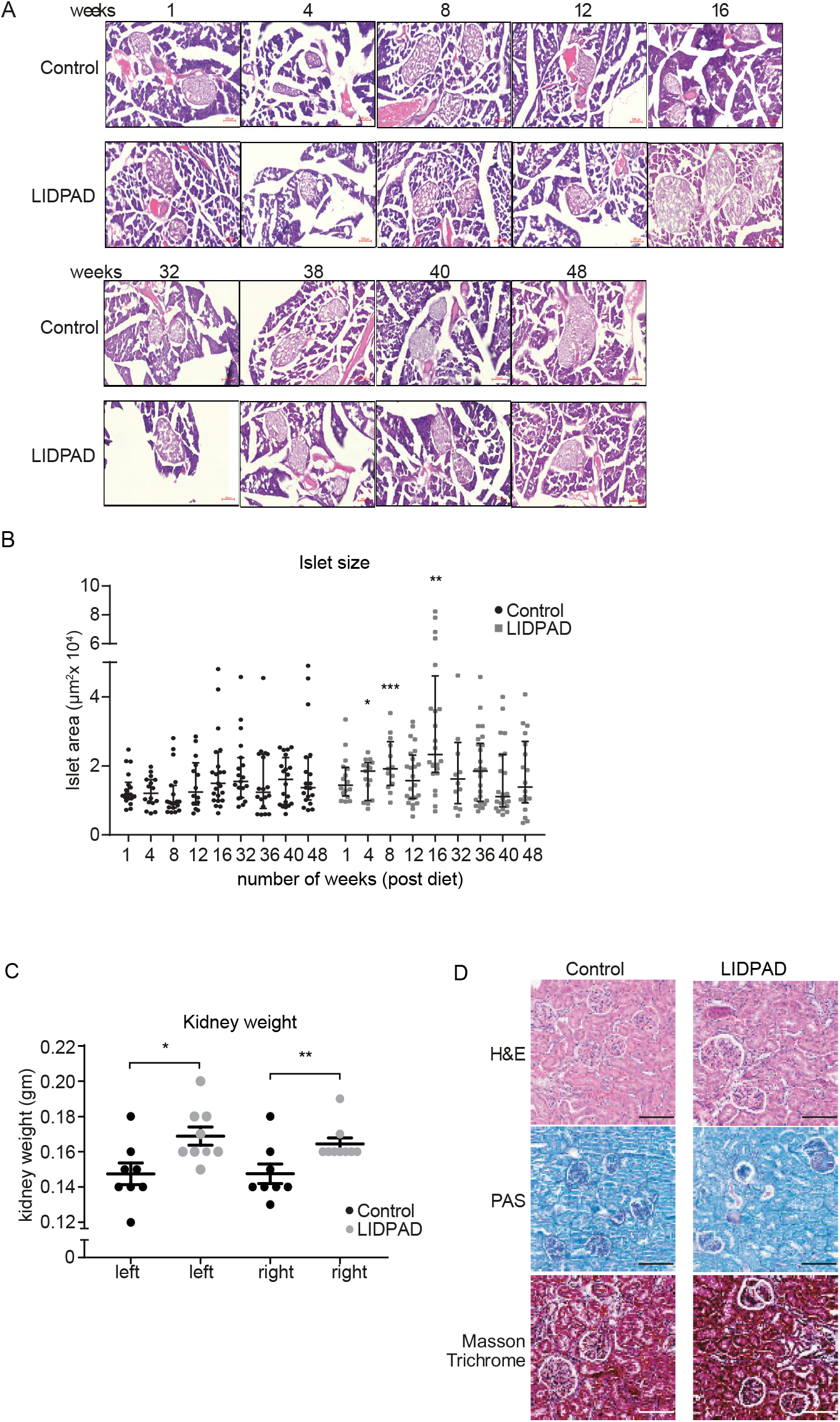
Histological images of pancreatic islets and kidneys from control and LIDPAD mice. **A.** Representative H&E images of pancreatic islets. **B.** Quantification and analysis of islet area using ImageJ for control and LIDPAD mice. across weeks 1 to 48. n = 12 - 28 region of interest (≥5 islets/animal). Data are expressed as the median ± interquartile range. **C.** Kidney weight (both kidneys) of control and LIDPAD mice compared at week 48. Data information: n = 8 per group. Data are expressed as the means ± SEMs, ***p<0.001, **p<0.01, *p<0.05 (Welch’s t test). n.s. denotes not significant. **D.** Representative images of kidneys stained using H&E to show the general morphology. Periodic Acid-Schiff (PAS) staining was used to highlight the basement membrane, and Masson trichrome staining was used to identify collagen deposition.

**Figure S4:**
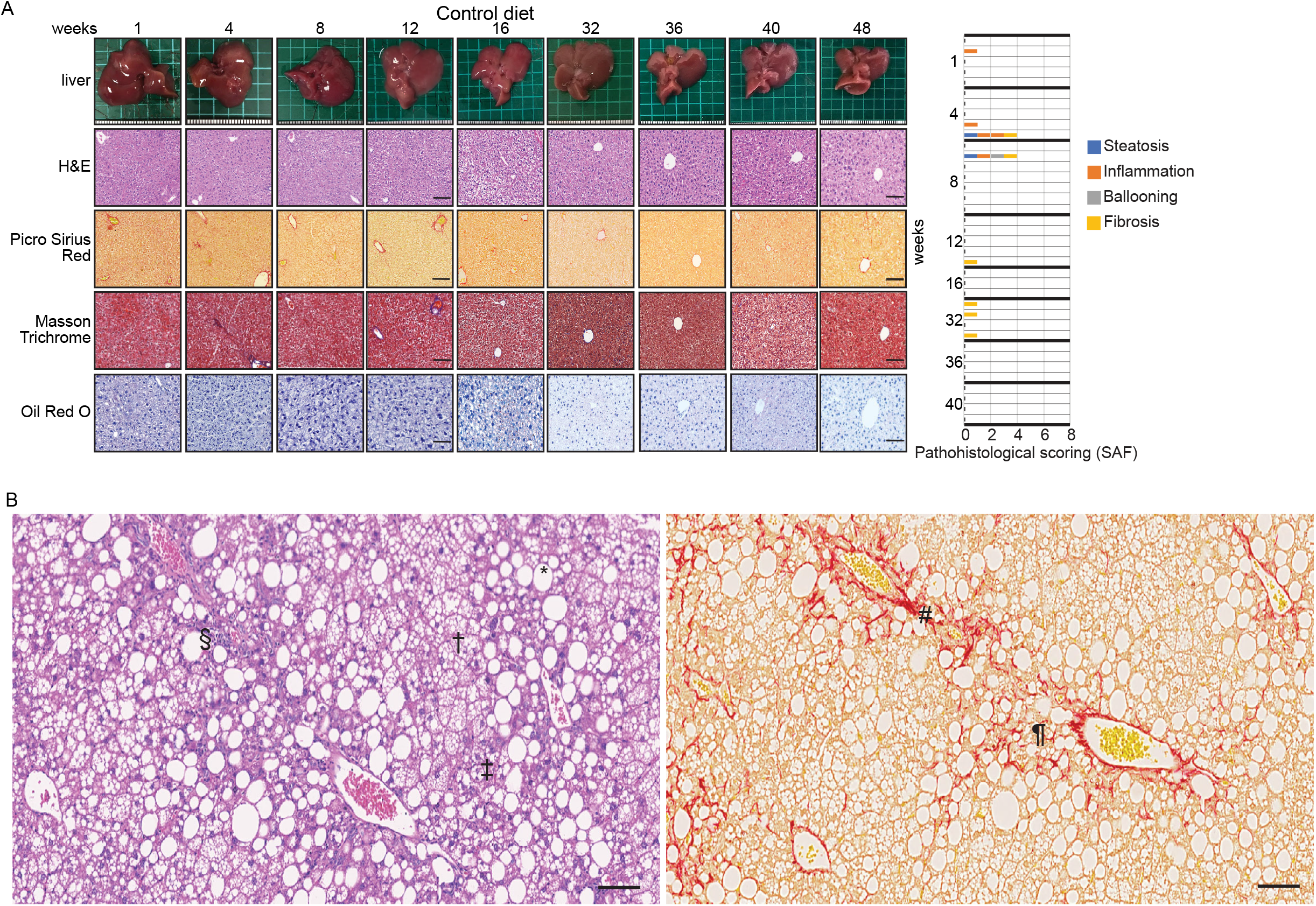
Representative macroscopic and microscopic images of the livers obtained from control and LIDPAD mice. **A.** Histological sections were stained with hematoxylin and eosin (H&E) to show general liver features, picrosirius red (PSR), and Masson’s trichrome (MT) to highlight collagen deposition, and oil red O (ORO) to detect the presence of lipids. The scale bar represents 100 μm. The graphs show the tabulated cumulative histological SAF scores of the livers from control mice at the indicated weeks postfeeding (right). Each row corresponds to one analysed liver. **B.** Representative histological images of mouse livers fed LIDPAD for 16 weeks, stained with H&E and PSR to indicate the hepatic pathological features. * indicates macrovesicular steatosis; † indicates microvesicular steatosis; ‡ indicates ballooned hepatocytes; § indicates the region of lobular inflammation; ¶ indicates the region of pericellular fibrosis; # indicates periportal fibrosis. The scale bar represents 100 μm.

**Figure S5:**
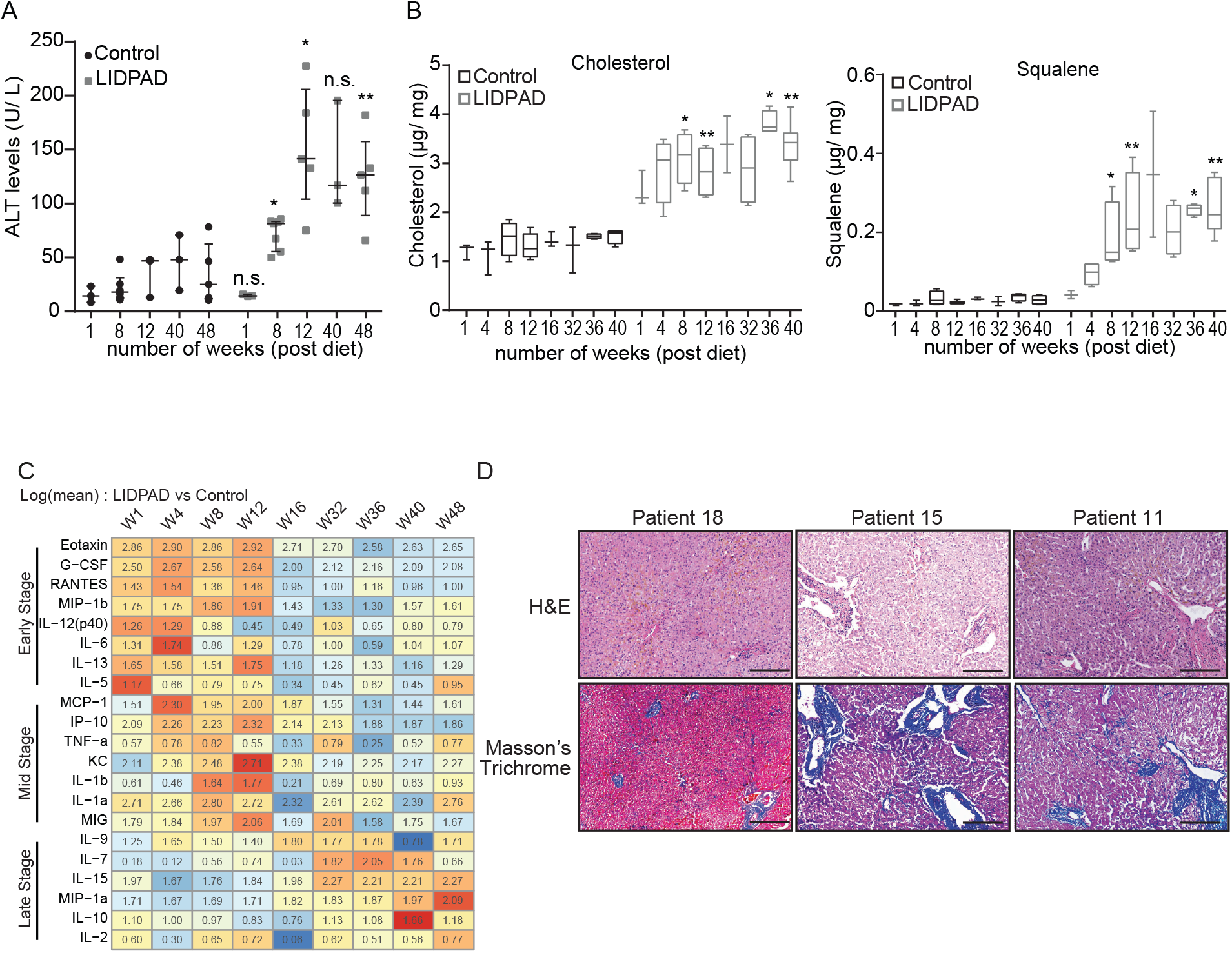
Hepatic ALT and serum cytokine levels. **A.** Blood chemistry readout of control and LIDPAD mice based on a liver function test (LFT) consisting of alanine transaminase (ALT). Data are median ± interquartile range. **B.** Lipid analysis highlighting cholesterol and squalene differences between control and LIDPAD mice. **C.** Heatmap of the serum cytokine array with color based on the row values after LIDPAD feeding for 1 to 48 weeks. Data are expressed as log (fold change of mean concentration) between LIDPAD and control mice. **D.** Representative images of liver biopsies from patients with NASH. Scale bar: 200 μm. For A-C: n = 3-7 per group. **p<0.01, *p<0.05 (2-way ANOVA with Sidak’s multiple comparison for C and Welch’s t test for D). n.s. denotes not significant.

**Figure S6:**
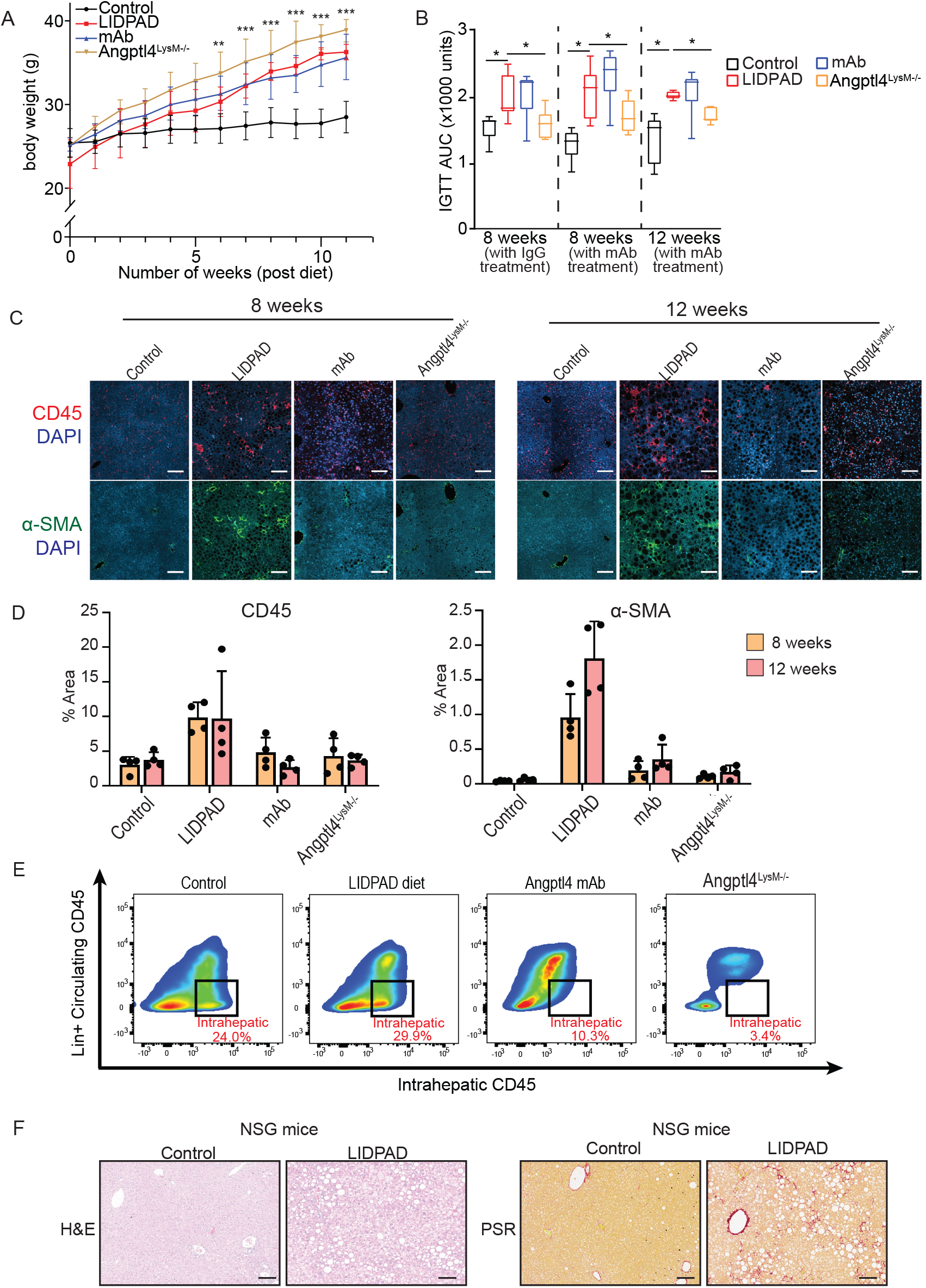
Metabolic effects of Angptl4 neutralizing antibodies on NASH mice. **A-B.** Body weight **(A)** and IGTT AUC **(B)** of mice in different treatment groups. AUC: area under the curve; IGTT, intraperitoneal glucose tolerance test. **C-D.** Representative immunofluorescence images **(C)** of livers from the indicated treatment groups after feeding for 8 and 12 weeks. CD45 indicates immune infiltration, while α-SMA indicates fibroblast activation. The scale bar represents 100 μm. Barplots showing the quantification of CD45 and α-SMA fluorescence intensities **(D)** in the immunofluorescence sections. **E.** FACS analysis showing the proportion of intrahepatic CD45+ immune cells in the indicated groups. **F.** Representative liver histological images of immunodeficient NSG mice fed a LIPDAD diet for 52 weeks. Data are expressed as the mean ± SD. *p<0.05, ***p<0.001. n= 6-9 mice per group.

**Figure S7:**
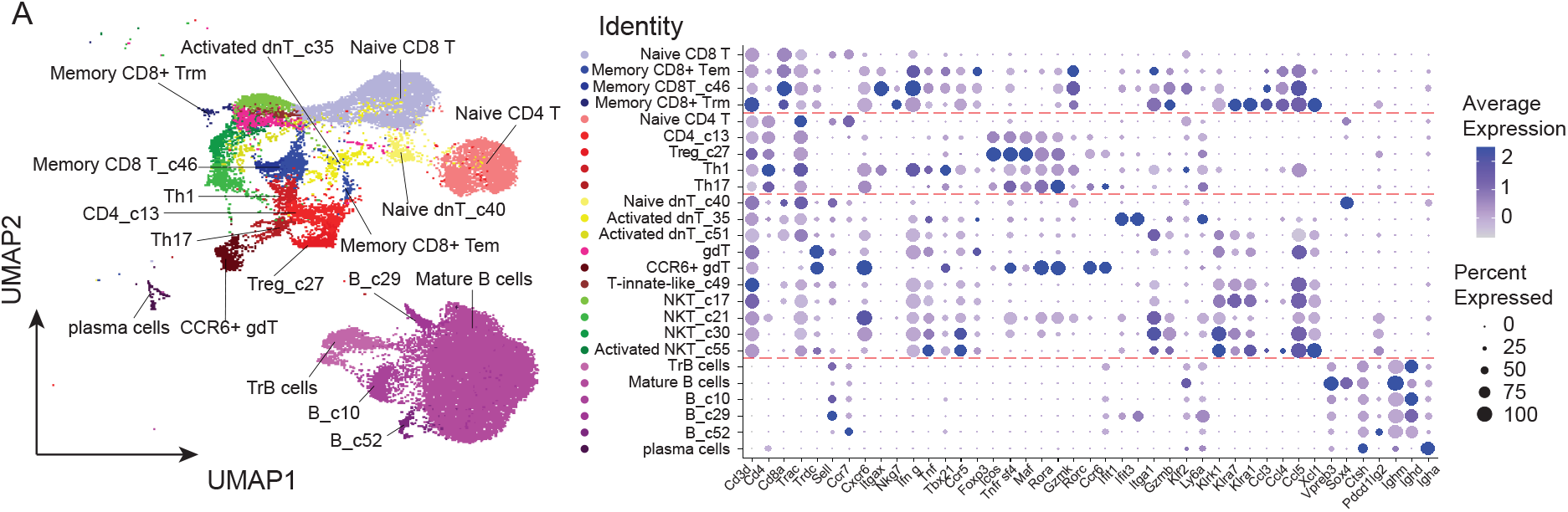
Intrahepatic B- and T-cell profiles. **A.** UMAP of adaptive immune subpopulations from B and T cells (left). Dot plot representing molecular markers used to identify specific subpopulations of B- and T cells (right). **B.** GSEA showing highly enriched gene sets in CD8+ T cells during NASH progression from 8 to 12 weeks of feeding based on the differential transcriptomes from different treatment groups. The normalized enrichment score (NES) is color-coded, where NES > 0 (red) and < 0 (blue) indicate gene functions that were enriched in up- and downregulated genes, respectively. Dot size denotes the −log10 FDR of representative enrichment tests.

**Figure S8:**
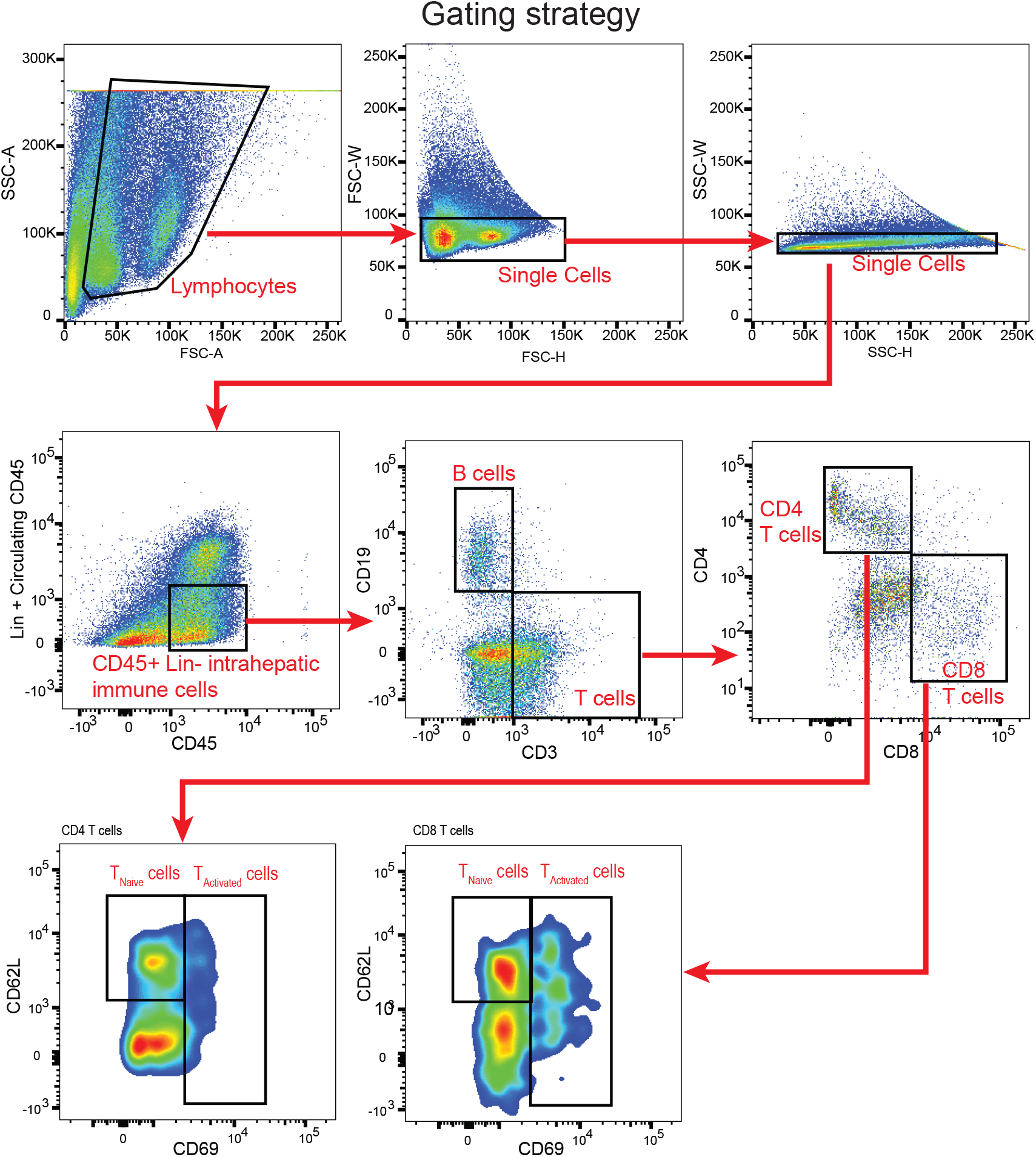
Gating strategy of FACS analysis to identify intrahepatic naïve and activated CD4^+^ and CD8^+^ T cells.

**Figure S9:**
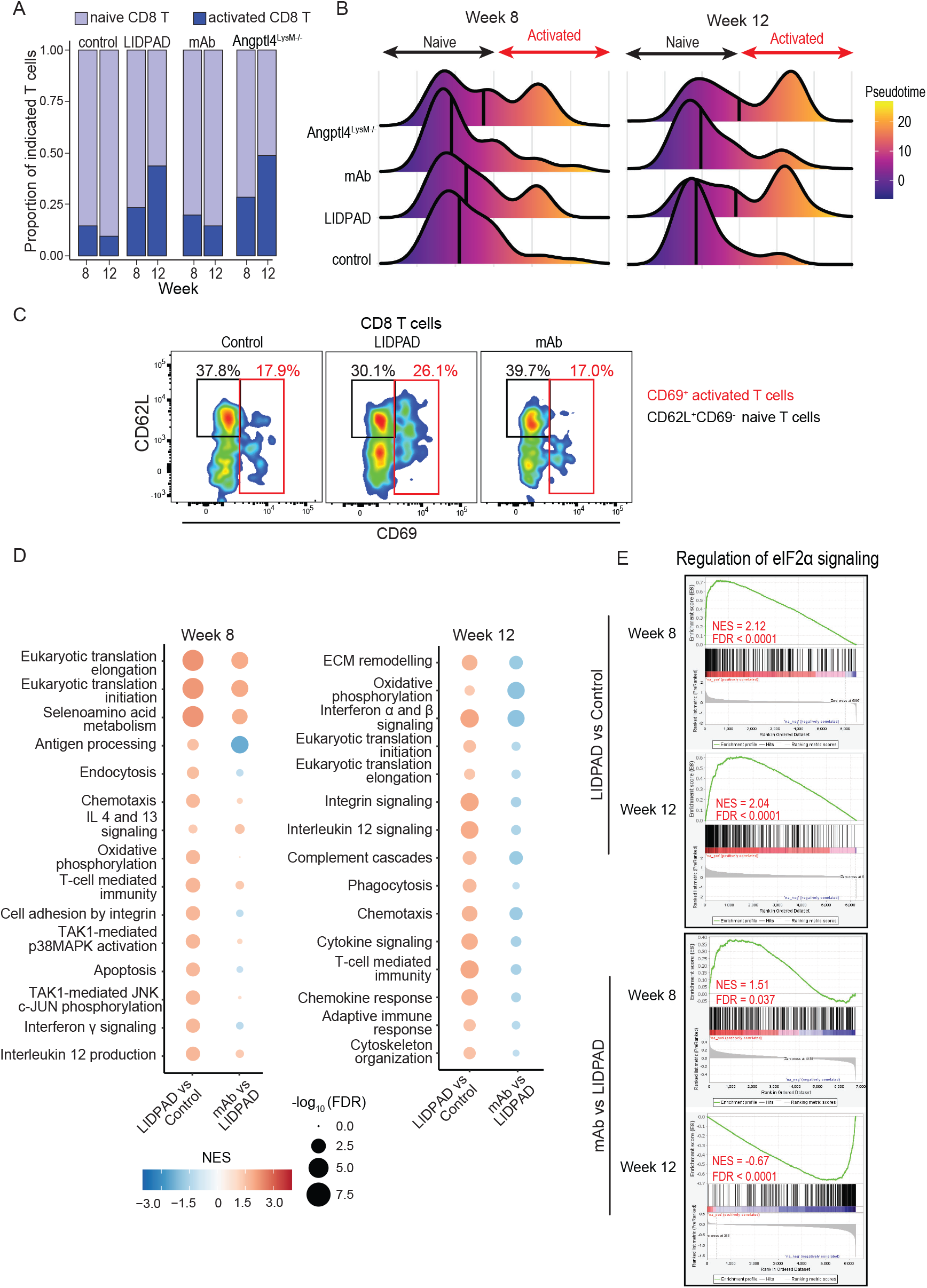
cAngptl4 mAb suppressed intrahepatic CD8^+^ T-cell activation. **A.** Proportion of intrahepatic naïve and activated CD8 T cells in control mice, LIDPAD-fed mice, and LIDPAD-fed mice treated with neutralizing monoclonal antibodies against cAngptl4 (mAb) after 8 and 12 weeks of feeding. **B.** Pseudotime analysis ordering the CD8 T cells according to the activation status. Vertical lines indicate the median pseudotime score of each group. **C.** Percentage of intrahepatic naïve (CD45+Lin-CD3+CD8+CD62L+CD69-) and activated (CD45+Lin-CD3+CD8+CD69+) CD8 T cells from the indicated groups after 12 weeks of feeding based on FACS. Lineage markers (Lin) include CD11b, NK1.1 and Ly6G. **D-E.** GSEA showing highly enriched gene sets in CD8+ T cells during NASH progression from 8 to 12 weeks of feeding **(D)** and signature genes involved in the regulation of eIF2α phosphorylation **(E)** based on the differential transcriptomes of naïve CD8 T cells from different treatment groups. Normalized enrichment score (NES) > 0 and < 0 indicate the enrichment of the gene set by up- and downregulated genes, respectively. For 5D, the NES is color-coded, and the dot size denotes the −log10 FDR of representative enrichment tests.

## Supplementary methods

### In Vivo Magnetic Resonance Imaging

Prior to in vivo imaging, animals were initially anaesthetized with 2.5-3 % isoflurane in combination with medical oxygen and medical air in a dedicated mouse chamber. Isoflurane concentration was reduced following induction to 1-2% during imaging to maintain respiration between 80-90 cycles/minute. Animal’s respiration rate and body temperature were monitored through a physiological monitoring system. Respiratory-linked gating with a 50 ms trigger delay was used during imaging. In vivo magnetic resonance imaging (Bruker 9.4 T Biospec) was performed using a 40 mm transmit/receive body coil. High-resolution anatomical T1-weighted images were acquired by a Fast Low Angle Shot (FLASH) sequence with repetition time (TR)-337 ms, echo time (TE) 2.5 ms, flip angle (FA) 30°, averages(AV)-3, image matrix size 256 x 256 and field of view (FOV)-40 x 40 mm. High-resolution anatomical T2 weighted images were acquired using RApid imaging with Refocused Echoes (RARE) sequence with TR-5175 ms, TE-30 ms, FA-90°, AV-2, image matrix size 256 × 256 and FOV-40 × 40 mm. Hepatic proton density fat fraction imaging was performed on LIDPAD (L) and control (C) mice at week 4 (n: L=3, C=3), week 8 (n: L=5, C=4) and week 16 (n, L=4, C=6) using multi-echo Dixon (m-Dixon) pulse sequence with parameters: TR-12 ms, TEs-1.85, 2.08, 2.32, 2.56, 2.80, 3.04, 3.27, 3.51 ms, FA-5°, slices-30, slice thickness-1 mm, images matrix size −256 × 256 and FOV-40 × 40 mm. The m-Dixon imaging data was processed and liver PDFF images were generated using the Fat-Water Toolbox. Identical regions of interest (ROI) were drawn within the liver of all animals by matching anatomical positions. Liver PDFF values were expressed as average percentages and statistical analysis (Mann–Whitney tests) was performed using SPSS software V23.

### Histological scoring of liver sections

The steatosis score (S) ranged from 0 to 3 (S0: <5%; S1: 5%-33%, mild; S2: 34%-66%, moderate; S3; >67%, marked). Activity grade (A, from 0-4) is the combination of hepatocyte ballooning (0-2) and lobular inflammation (0-2). Ballooning was scored as 0 (normal polygonal hepatocytes), 1 (round, not enlarged hepatocytes, reticulated cytoplasm), and 2 (2x enlarged hepatocytes with clear cytoplasm and clumping of intermediate fibers). Lobular inflammation was scored as 0 (none), 1 (≤2 foci per 20x field), and 2 (>2 foci/per 20x field). Fibrosis scoring was as follows: stage 0 (F0): none; stage 1 (F1): 1a or 1b delicate or dense zone 3 perisinusoidal fibrosis, respectively, 1c periportal fibrosis only; stage 2 (F2): zone 3 perisinusoidal fibrosis and periportal fibrosis; stage 3 (F3): bridging fibrosis; and stage 4 (F4): cirrhosis. Portal inflammation was noted as absent or present.

### Histological preparation of kidney samples

Kidneys were fixed in buffered formaldehyde solution, processed, and embedded in paraffin. Four-micrometer-thick tissue sections were stained with H&E, Periodic Acid-Schiff (PAS) Stain Kit (ab150680), and MT stain, according to the manufacturer’s protocol. All slide images were captured using a Carl Zeiss Axio Slide Scanner Z1.

### Pancreas harvesting, H&E staining, and islet size determination

Harvested pancreatic specimens were fixed in 4% paraformaldehyde (PFA) for 16 h followed by cryoprotection in 30% sucrose for an additional 24 h. The specimens were embedded and sectioned in Optimal Cutting Temperature compound (OCT, Sakura, Japan) and stained with H&E. The stained slides were imaged using a Carl Zeiss Axio Slide Scanner. Z1 at 20X magnification. Zoomed-in-tif images of the islets were documented together with a digital ruler and annotations. Islets were marked and analysed for size using ImageJ software. The scale was adjusted for each image, and the ROI for each islet, manually marked, was used to calculate the area. Islet areas were then collated and evaluated on GraphPad Prism. Data are presented as the means ± SEMs (≥5 islets/animal).

### Insulin tolerance test

Mice were injected intraperitoneally (i.p.) with 1 U/kg body weight insulin (Lilly, USA) after 4 h of fasting. Blood glucose concentrations were measured up to 120 min after injection from the tail vein using a glucometer (Accucheck Performa, Roche, USA). All measurements were taken from 10 am-12 noon the next day.

### Intraperitoneal Glucose Tolerance Test

Mice were injected intraperitoneally with 2.0 g/kg body weight glucose after 16 h of overnight fasting. Blood glucose concentrations were measured up to 120 min after injection from the tail vein using a glucometer (Accucheck Performa, Roche, USA). All measurements were taken from 10 am-12 noon the next day.

### Serum cytokine array

Blood samples collected from the mice via cardiac were left to clot for 30 minutes at room temperature and then transferred to ice. The clotted samples were spun in the centrifuge at 12,000 × g for 1 min to separate the serum. The supernatant was carefully retrieved and stored. A mouse cytokine 32-plex discovery assay was then performed by Eve Technologies (Calgary, Canada). *Calorimetric chamber*. Mice were prehabituated in the testing room at 30 °C for 5 days, including for 3 days in habituation cages. The climate chamber was maintained at a constant temperature of 30 °C and humidity of 50%. The light was ON at 7 am with 80 lux in the chamber and OFF at 7 pm. Animals had ad libitum access to food (LIDPAD or control accordingly) and filtered water.

Oxygen and carbon dioxide sensors were calibrated with calibration gas mixtures (calibration report: good air equivalent: CO_2_: 0.050 ± 0.0005%, O_2_: 20.900 ± 0.1% in N2; CO_2_ span: CO_2_: 1.000 ± 0.001%, O_2_: 20.000 ± 0.1% in N2). The sample air flow was adjusted to 0.37 L/min. High-precision weighing stations combined with leak- and spill-proof containers recorded the body weight, food intake, and water intake. Spontaneous activity was recorded with two levels of infrared light beam frames surrounding each cage. Recording began from the chamber’s first entrance, but measures were considered only from the 2nd day in the chamber. Running wheels were added to the cage from the 5th day. Habituation with running wheels lasted 2.5 days before the data were considered. Measurements were taken every 15 min and are presented here hourly or as the means per hour.

### Lipid analyses using liquid chromatography–mass spectrometry

Lipids were extracted from mouse liver tissue using liquid–liquid extraction as described(*1*) with modifications. Briefly, 10-20 mg of tissue was mechanically homogenized in a methanol:MTBE mixture using bead beating (Precellys). The lysates were further subjected to sonication in an ultrasonic bath with ice for 10 min. Phase separation was induced by adding water, followed by a 10 min incubation at room temperature. The sample was then centrifuged at 1,000 × g for 10 min. The upper organic phase was collected, and the aqueous phase was re-extracted. The pooled organic phase was dried under vacuum using a Speedvac, and the lipid extracts were stored at −80 °C prior to further analyses.

Samples were spiked with internal standards containing d7-cholesterol (Avanti Lipids), and sterols were analysed using atmospheric pressure chemical ionization (APCI) triple quadrupole mass spectrometry (SCIEX Qtrap 6500+) with upfront liquid chromatography (Agilent Technologies). The separation of sterols was achieved using a Poroshell 120 SB-C18 column (3.0 × 50 mm, 2.7 μM, Agilent Technologies), which is a modification of a previous method (*2*). For mass spectrometry analyses, the instrument was operated in multiple reaction mode (MRM). For quantitation, the area under the curve was obtained for each sterol measured and normalized to the internal standard and tissue weight. External calibration was performed for cholesterol and squalene to adjust the response factor for squalene relative to cholesterol.

### Immunofluorescence staining

Liver sections were dewaxed, rehydrated and subject to heat antigen retrieval in sodium citrate buffer (10 mM, 0.05% Tween-20, pH 6) using Aptum Biologics 2100 Antigen Retriever (Aptum Biologics, UK). The sections were blocked with 5% (v/v) fetal bovine serum for an hour and labelled with anti-mouse CD45 or anti-mouse α-smooth muscle actin (α-SMA) antibodies (Cell Signaling Technology, USA) at 4°C overnight. The sections were then treated with Alexa Fluor 680-conjugated secondary antibodies and counterstained with Hoechst 33342. Microscopic images of the sections were captured with Zeiss AxioScan.Z1.

### Single-cell RNA sequencing (scRNA-seq)

Single-cell droplets and cDNA libraries were prepared using a Chromium Single Cell 3’ V3 kit (10X Genomics, USA) according to the manufacturer’s instructions. The cDNA libraries were sequenced on a NovaSeq6000 (Illumina, USA). Raw sequencing reads were aligned to the mm10 mouse reference genome using Cell Ranger (v3.1.0) to generate single-cell count matrices that were normalized, integrated, and annotated using Seurat (v3.0)(*3*). The obtained count matrices were merged and normalized using SCTransform. Subsequently, cells with fewer than 250 genes, more than 20% mitochondrial genes, and genes with less than 500 UMI counts were filtered out. Average housekeeping gene counts were used as a quality control measure. Datasets were not corrected for cell cycle differences, as visual inspection of the first 2 principal components of cell cycle genes did not show cell cycle bias. Overall, a total of 32551 cells were obtained across all 8 conditions. Subsequently, the dataset is integrated using the Seurat v4 pipeline. The kBET metric revealed the effective removal of the batch effect. Principal component analysis (PCA) was performed on highly variable genes, and the first 35 PCA components were used to construct UMAP. Unsupervised cell clustering was performed with the Louvain algorithm. Cluster marker detection was performed by differentially expressed gene (DEG) analysis for each marker against the remaining markers using FindAllMarkers (Supplementary file). Functional annotation of differentially expressed gene sets was performed using EnrichR(*4*). Trajectory and pseudotime DEG analyses were performed using Slingshot(*5*) and Trade-seq(*6*). The single-cell sequencing data were deposited in GEO (Accession number: GSE214172).

### Annotation of the scRNAseq immune subpopulation

Cell type annotation was performed automatically using the SingleR package(*7*) against the ImmGen(*8*) database, as it performed superior in a recent benchmarking evaluation(*9*). Subsequently, manual curation of cell types was performed with the top differentially expressed genes of each cluster and compared to automatic annotation to ensure fidelity of annotation(10).

T lymphocytes are identified with expression of canonical Cd3 marker – where it is segregated into alpha-beta T-cell receptor (TCR) T lymphocytes and gd-T cells (expressing gamma-delta TCR) via Trac or Trdc markers, respectively. Furthermore, conventional CD4^+^ and CD8^+^ T cells were identified with Cd4, Cd8a, and Cd8b1 markers. Naïve T cells were identified with L-selectin (Sell) within these populations. Cells that ambiguously expressed CD4 and CD8 markers were identified as double-negative T cells (dnT). T-innate-like and NKT-cell types are enriched with innate-like markers such as Nkg7 and the Klr gene family. Murine intrahepatic NKT cells specifically express the Cxcr6 marker, which indicates the ability to recognize Cd1d+ antigen presenting cells(*11*).

B lymphocytes were identified with enrichment of canonical Ms4a1, Cd19, and Cd79a markers. B lymphocytes also express H2-Aa, which are major histocompatibility class 2 (MHC-II) molecules (*12*) that are functionally involved in antigen presentation to CD4+ T cells during activation. Antibody-producing plasma cells were identified with the Igha marker. Innate lymphoid cells (ILCs) are differentiated from NK cells with high expression of Xcl1 and Cxcr6 and lower expression of Nkg7(*13*). Neutrophils were identified with S100a8 and S100a9 markers(*11*). Others have reported technical dropout issues in scRNA-seq to identify neutrophils with canonical Ly6g mRNA markers(*13*). Monocytes express Fcgr1, Fcgr3, and Mafb and lack the Itgax activation marker. Kupffer cells are differentiated from monocyte-derived macrophages (mo-macs) with exclusive expression of Clec4f+ and Csf1r+(*14*). Finally, classical dendritic cells (cDCs) were identified with Clec10a and H2-Aa, while plasmacytoid DCs (pDCs) were identified with Tcf4 and Irf8 markers(*15, 16*).

