## Supplemental Information for "Angiopoietin-like 4 shapes the intrahepatic T-cell landscape via eIF2α signaling during steatohepatitis in diet-induced NAFLD"

A

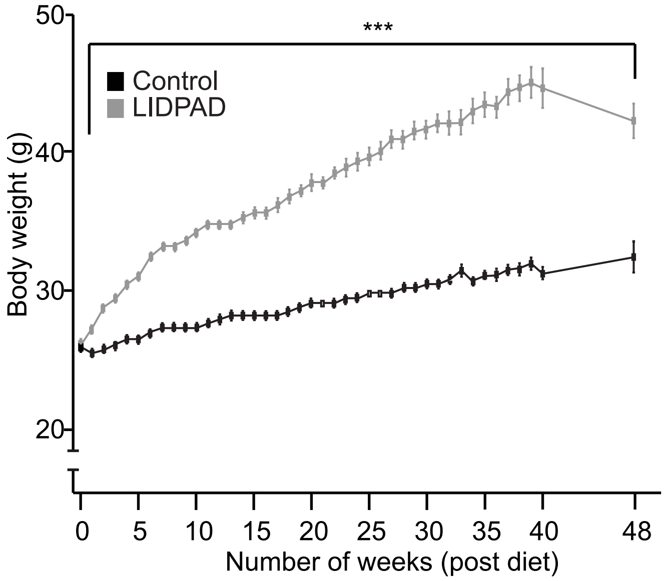


| Duration on diet | Number of animals | |
|---|---|---|
| | Control | LIDPAD |
| 1 | 84 | 90 |
| 4 | 64 | 71 |
| 8 | 51 | 57 |
| 12 | 38 | 47 |
| 16 | 33 | 37 |
| 32 | 30 | 34 |
| 36 | 20 | 25 |
| 40 | 13 | 17 |
| 48 | 8 | 9 |

B

IGTT Curves

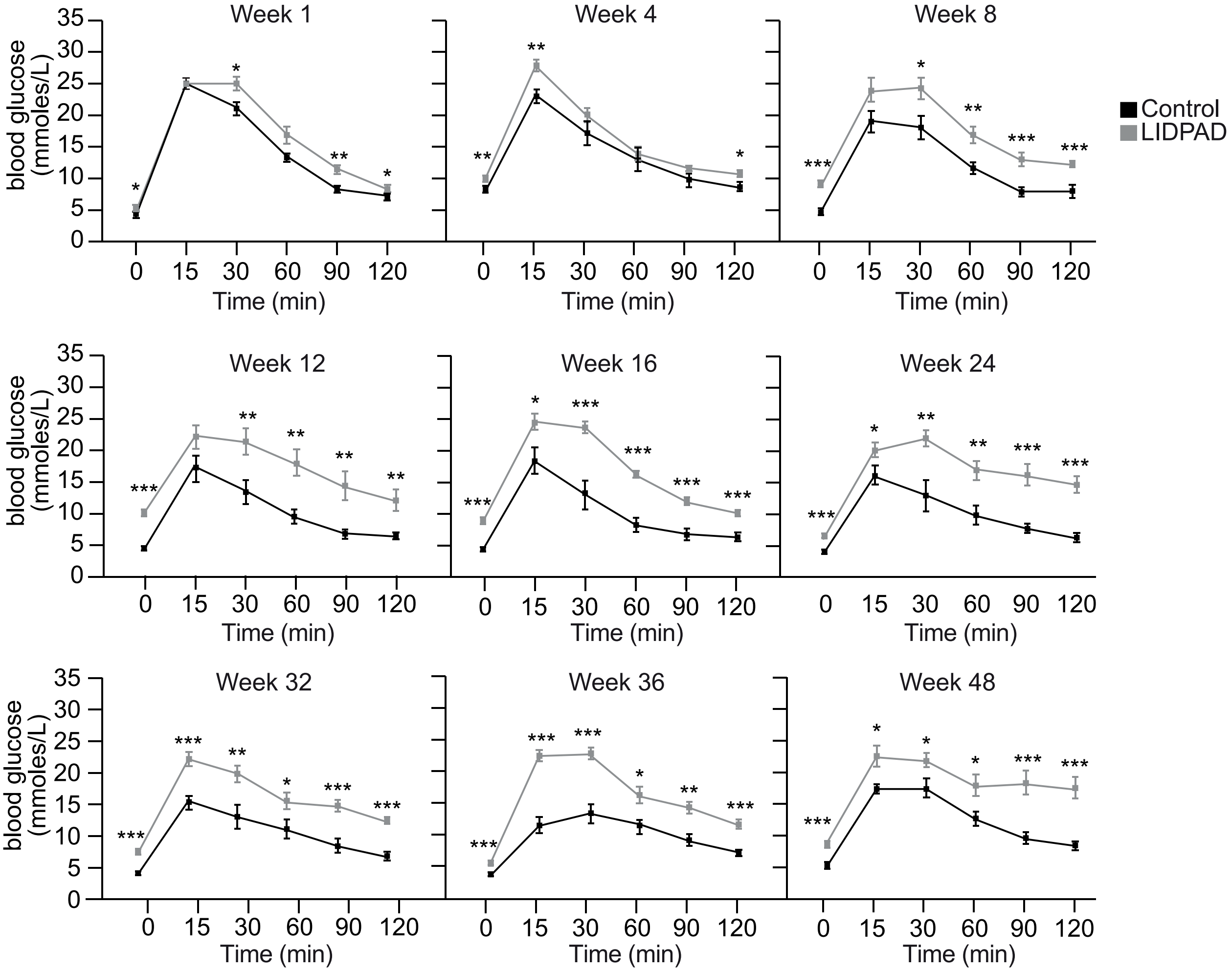

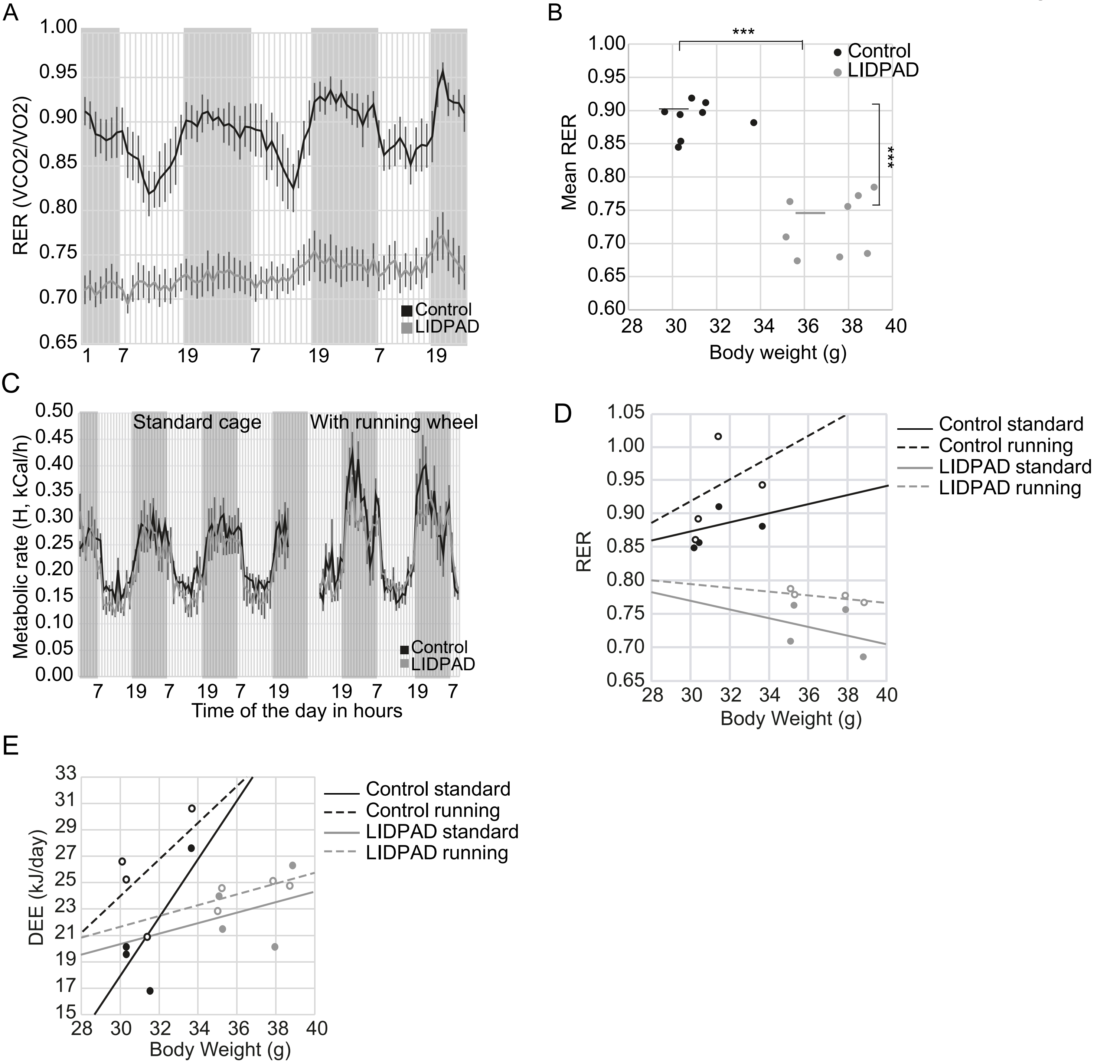
A
RER (VCO2/VO2)
1.00
0.95
0.90
0.85
0.80
0.75
0.70
0.65
1 7 19 7 19 7 19
Control
LIDPAD
B
Mean RER
***
Control
LIDPAD
Body weight (g)
28 30 32 34 36 38 40
C
Standard cage
With running wheel
Metabolic rate (H, kCal/h)
Time of the day in hours
Control
LIDPAD
D
RER
Control standard
Control running
LIDPAD standard
LIDPAD running
Body Weight (g)
E
DEE (kJ/day)
Control standard
Control running
LIDPAD standard
LIDPAD running
Body Weight (g)

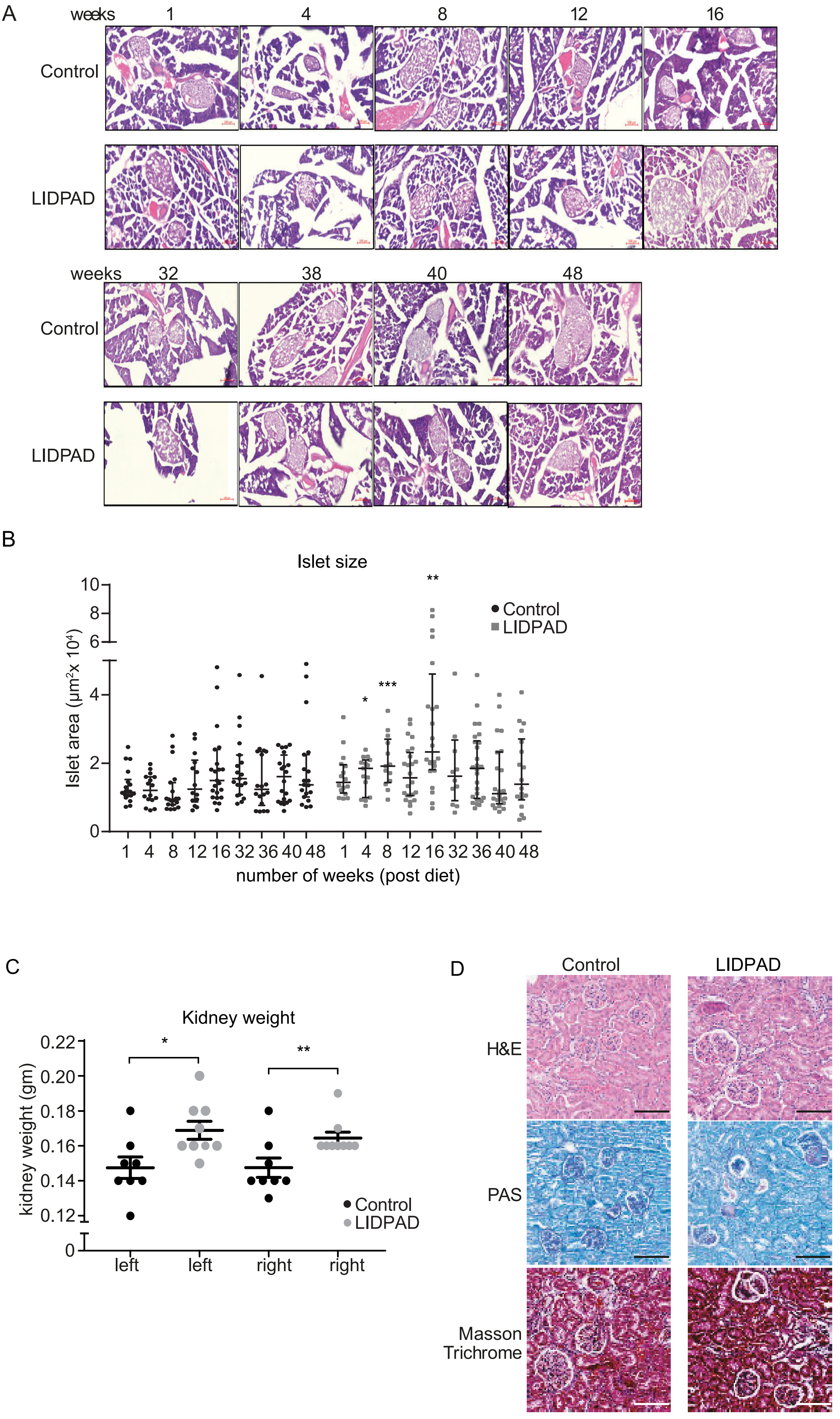
A
weeks
1
4
8
12
16
Control
LIDPAD
weeks
32
38
40
48
Control
LIDPAD
B
Islet size
Islet area (µm²x 10⁴)
10
8
6
4
2
0
**
*
***
Control
LIDPAD
1 4 8 12 16 32 36 40 48 1 4 8 12 16 32 36 40 48
number of weeks (post diet)
C
Kidney weight
kidney weight (gm)
0.22
0.20
0.18
0.16
0.14
0.12
0
*
**
Control
LIDPAD
left
left
right
right
D
Control
LIDPAD
H&E
PAS
Masson
Trichrome

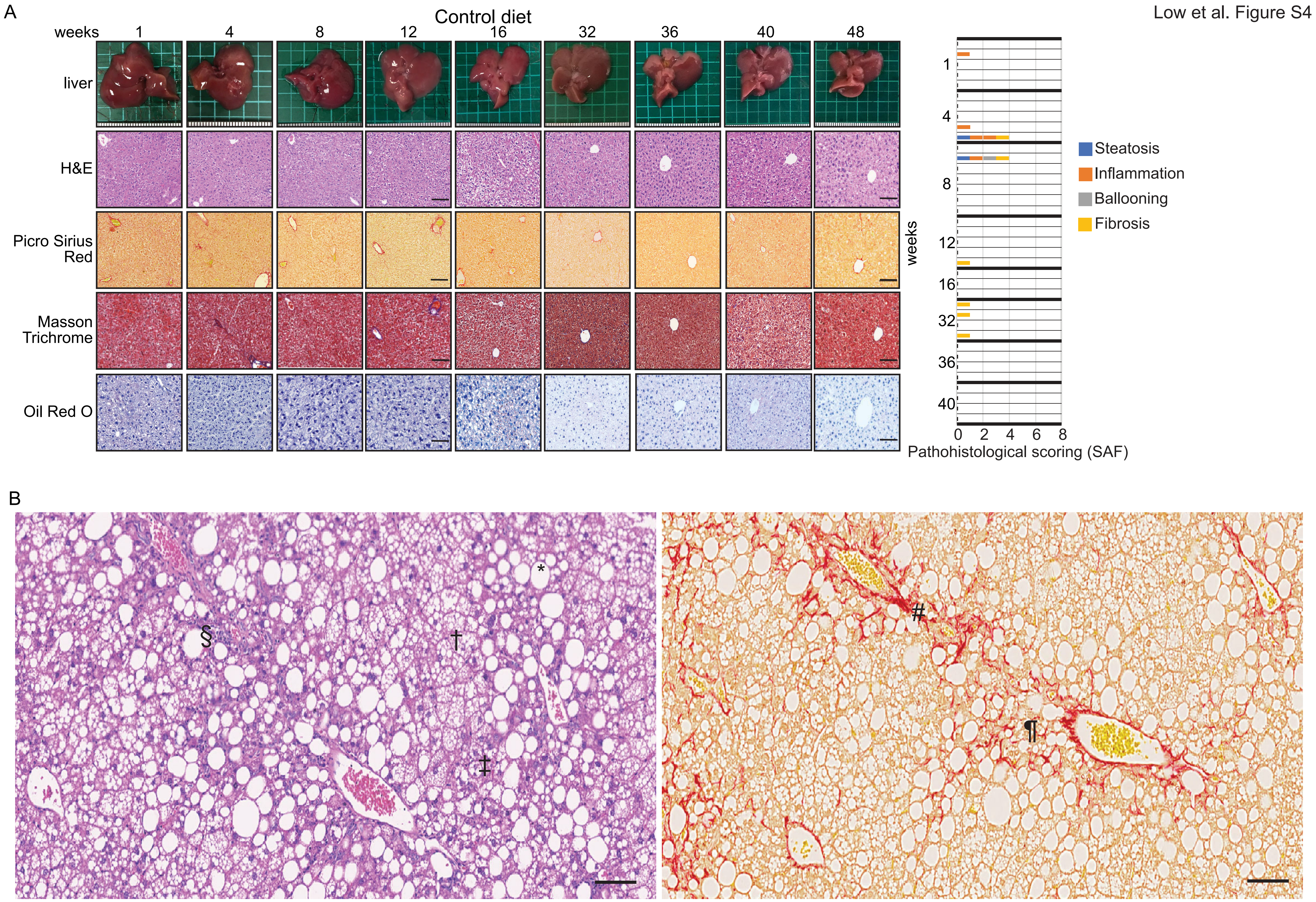

Low et al. Figure S4
A
Control diet
weeks
1
4
8
12
16
32
36
40
48
liver
H&E
Picro Sirius Red
Masson Trichrome
Oil Red O
Steatosis
Inflammation
Ballooning
Fibrosis
weeks
Pathohistological scoring (SAF)
0
2
4
6
8
B
§
†
*
‡
#
¶

A

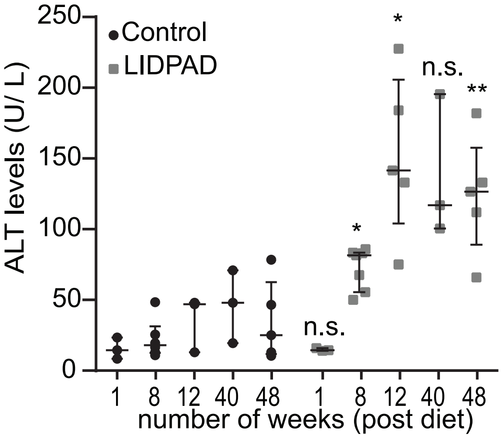


B

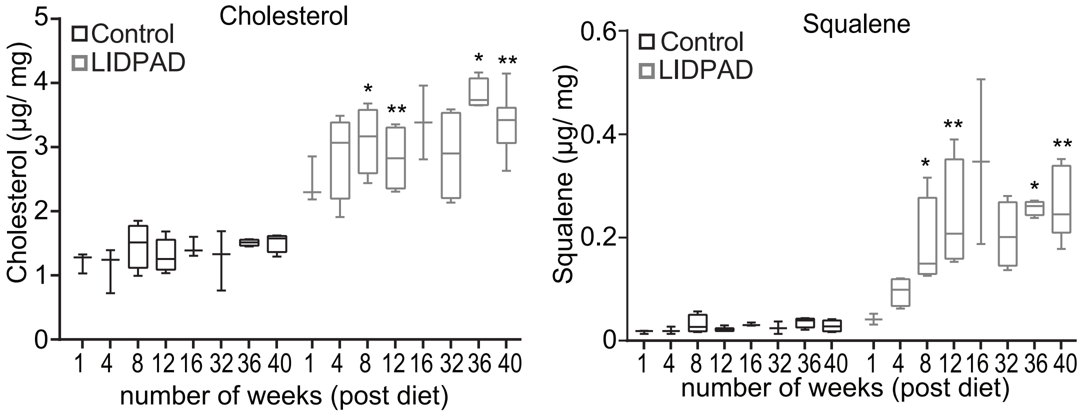


C

Log(mean) : LIDPAD vs Control

| Stage | | W1 | W4 | W8 | W12 | W16 | W32 | W36 | W40 | W48 |
|---|---|---|---|---|---|---|---|---|---|---|
| Early Stage | Eotaxin | 2.86 | 2.90 | 2.86 | 2.92 | 2.71 | 2.70 | 2.58 | 2.63 | 2.65 |
| | G−CSF | 2.50 | 2.67 | 2.58 | 2.64 | 2.00 | 2.12 | 2.16 | 2.09 | 2.08 |
| | RANTES | 1.43 | 1.54 | 1.36 | 1.46 | 0.95 | 1.00 | 1.16 | 0.96 | 1.00 |
| | MIP−1b | 1.75 | 1.75 | 1.86 | 1.91 | 1.43 | 1.33 | 1.30 | 1.57 | 1.61 |
| | IL−12(p40) | 1.26 | 1.29 | 0.88 | 0.45 | 0.49 | 1.03 | 0.65 | 0.80 | 0.79 |
| | IL−6 | 1.31 | 1.74 | 0.88 | 1.29 | 0.78 | 1.00 | 0.59 | 1.04 | 1.07 |
| | IL−13 | 1.65 | 1.58 | 1.51 | 1.75 | 1.18 | 1.26 | 1.33 | 1.16 | 1.29 |
| | IL−5 | 1.17 | 0.66 | 0.79 | 0.75 | 0.34 | 0.45 | 0.62 | 0.45 | 0.95 |
| Mid Stage | MCP−1 | 1.51 | 2.30 | 1.95 | 2.00 | 1.87 | 1.55 | 1.31 | 1.44 | 1.61 |
| | IP−10 | 2.09 | 2.26 | 2.23 | 2.32 | 2.14 | 2.13 | 1.88 | 1.87 | 1.86 |
| | TNF−a | 0.57 | 0.78 | 0.82 | 0.55 | 0.33 | 0.79 | 0.25 | 0.52 | 0.77 |
| | KC | 2.11 | 2.38 | 2.48 | 2.71 | 2.38 | 2.19 | 2.25 | 2.17 | 2.27 |
| | IL−1b | 0.61 | 0.46 | 1.64 | 1.77 | 0.21 | 0.69 | 0.80 | 0.63 | 0.93 |
| | IL−1a | 2.71 | 2.66 | 2.80 | 2.72 | 2.32 | 2.61 | 2.62 | 2.39 | 2.76 |
| | MIG | 1.79 | 1.84 | 1.97 | 2.06 | 1.69 | 2.01 | 1.58 | 1.75 | 1.67 |
| Late Stage | IL−9 | 1.25 | 1.65 | 1.50 | 1.40 | 1.80 | 1.77 | 1.78 | 0.78 | 1.71 |
| | IL−7 | 0.18 | 0.12 | 0.56 | 0.74 | 0.03 | 1.82 | 2.05 | 1.76 | 0.66 |
| | IL−15 | 1.97 | 1.67 | 1.76 | 1.84 | 1.98 | 2.27 | 2.21 | 2.21 | 2.27 |
| | MIP−1a | 1.71 | 1.67 | 1.69 | 1.71 | 1.82 | 1.83 | 1.87 | 1.97 | 2.09 |
| | IL−10 | 1.10 | 1.00 | 0.97 | 0.83 | 0.76 | 1.13 | 1.08 | 1.66 | 1.18 |
| | IL−2 | 0.60 | 0.30 | 0.65 | 0.72 | 0.06 | 0.62 | 0.51 | 0.56 | 0.77 |

D

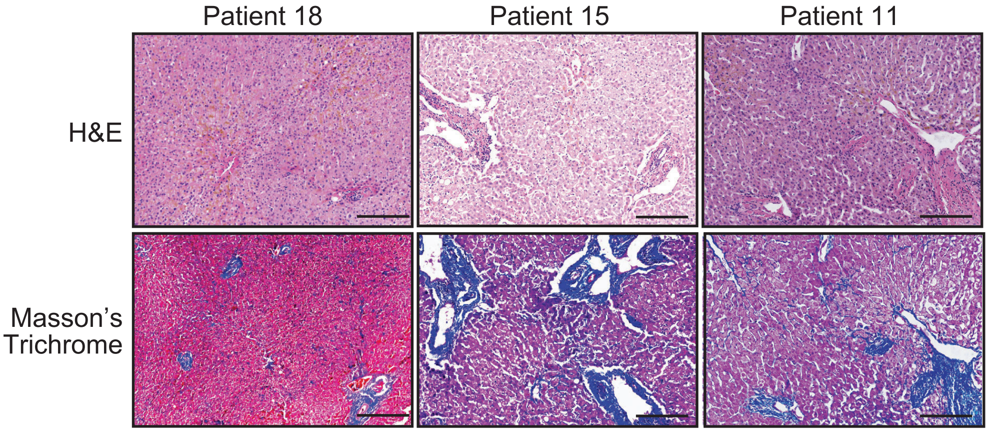

Low et al. Figure S6

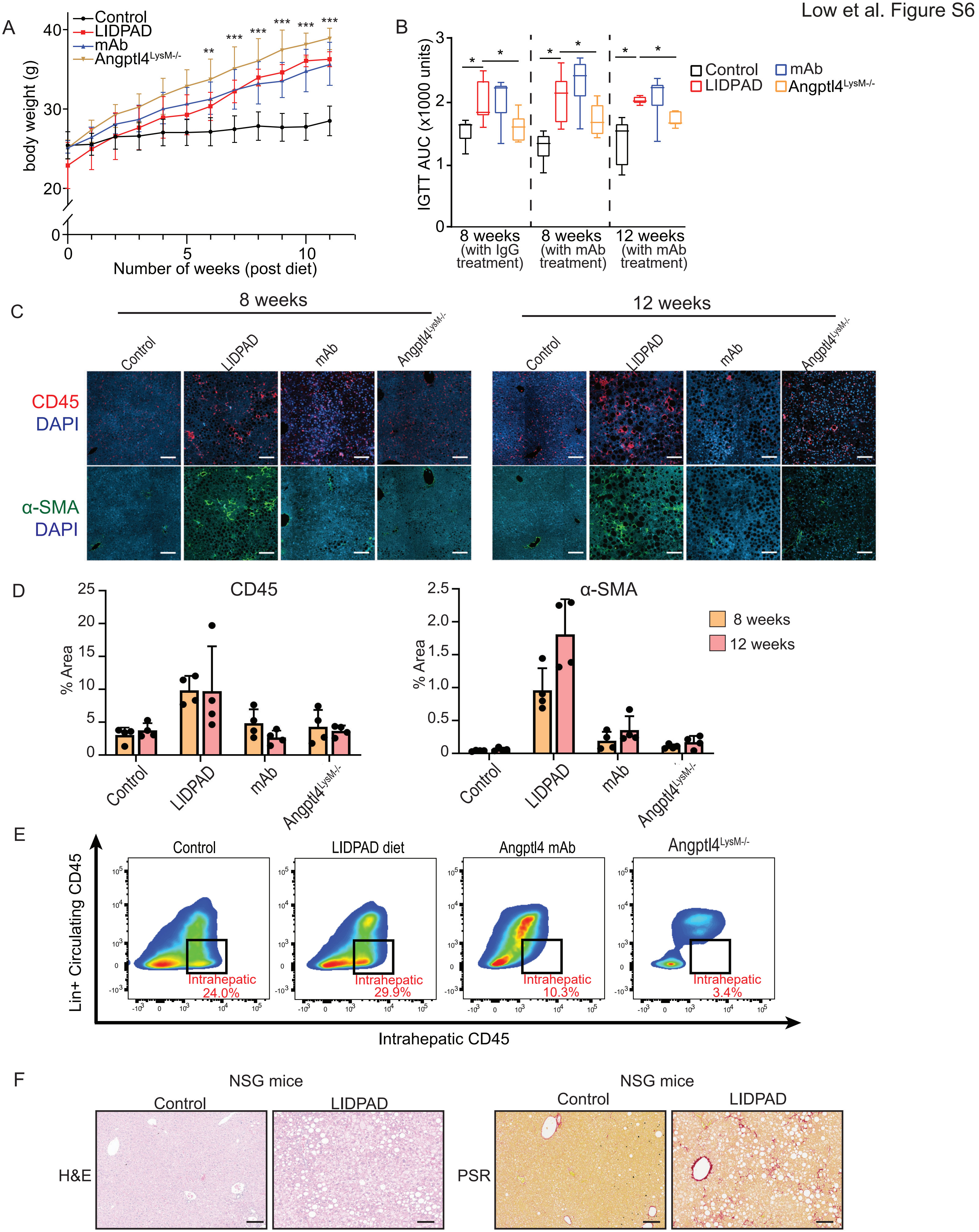

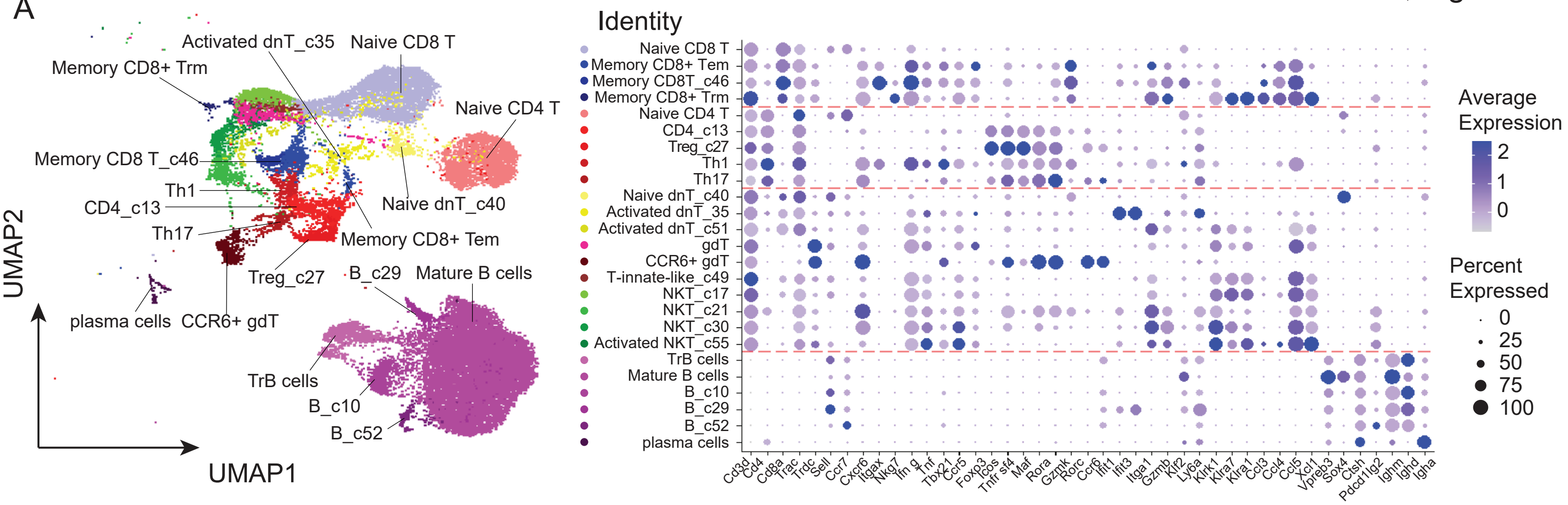
A
Activated dnT_c35
Naive CD8 T
Memory CD8+ Trm
Naive CD4 T
Memory CD8 T_c46
Th1
CD4_c13
Naive dnT_c40
Th17
Memory CD8+ Tem
Treg_c27
B_c29
Mature B cells
plasma cells
CCR6+ gdT
TrB cells
B_c10
B_c52
UMAP2
UMAP1
Identity
Naive CD8 T
Memory CD8+ Tem
Memory CD8T_c46
Memory CD8+ Trm
Naive CD4 T
CD4_c13
Treg_c27
Th1
Th17
Naive dnT_c40
Activated dnT_35
Activated dnT_c51
gdT
CCR6+ gdT
T-innate-like_c49
NKT_c17
NKT_c21
NKT_c30
Activated NKT_c55
TrB cells
Mature B cells
B_c10
B_c29
B_c52
plasma cells
Cd3d
Cd4
Cd8a
Trac
Trdc
Sell
Ccr7
Cxcr6
Itgax
Nkg7
Ifn g
Tnf
Tbx21
Ccr5
Foxp3
Icos
Tnfr sf4
Maf
Rora
Gzmk
Rorc
Ccr6
Ifit1
Ifit3
Itga1
Gzmb
Klf2
Ly6a
Klrk1
Klra7
Klra1
Ccl3
Ccl4
Ccl5
Xcl1
Vpreb3
Sox4
Ctsh
Pdcd1lg2
Ighm
Ighd
Igha
Average Expression
2
1
0
Percent Expressed
0
25
50
75
100

### Gating strategy

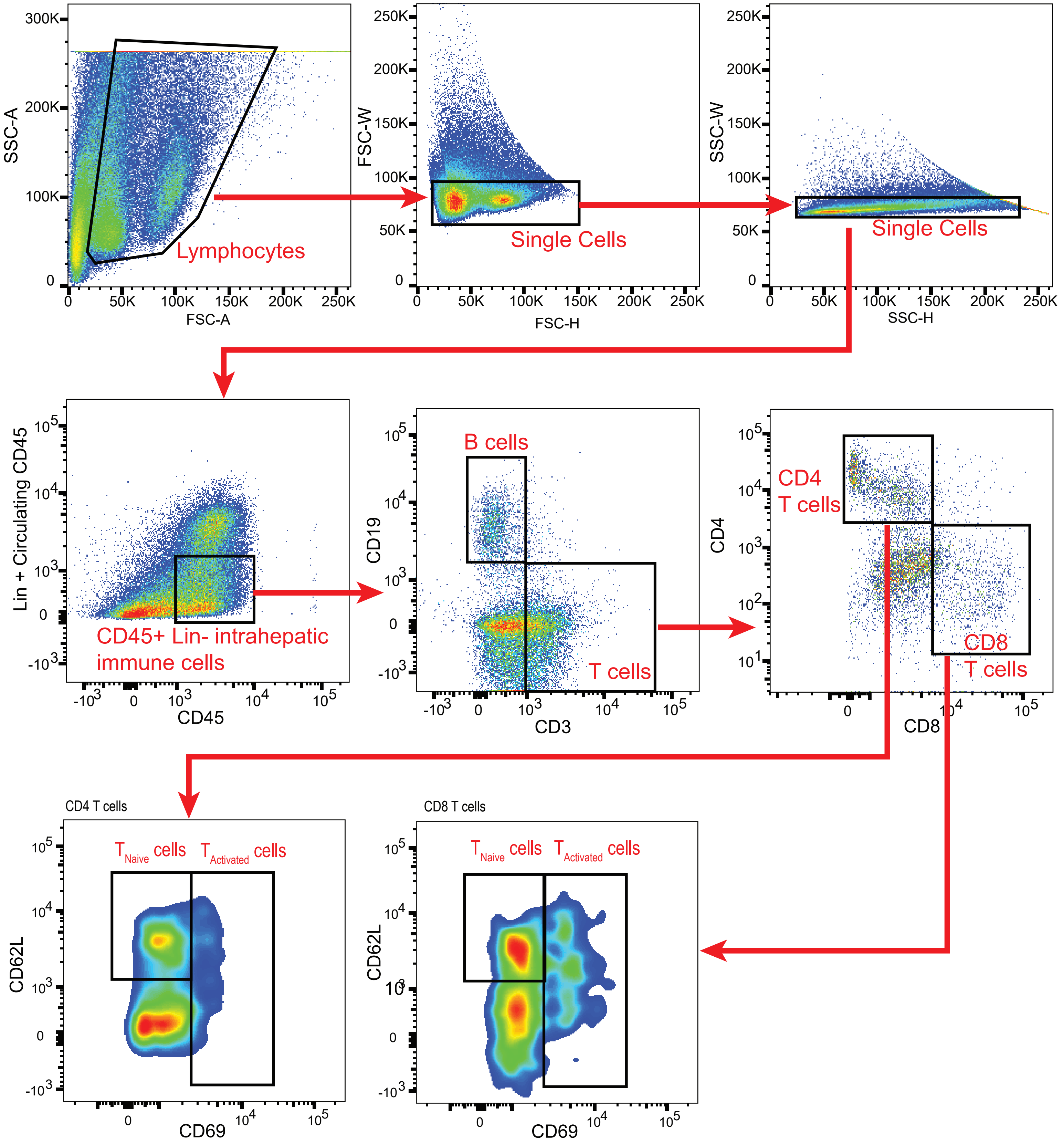

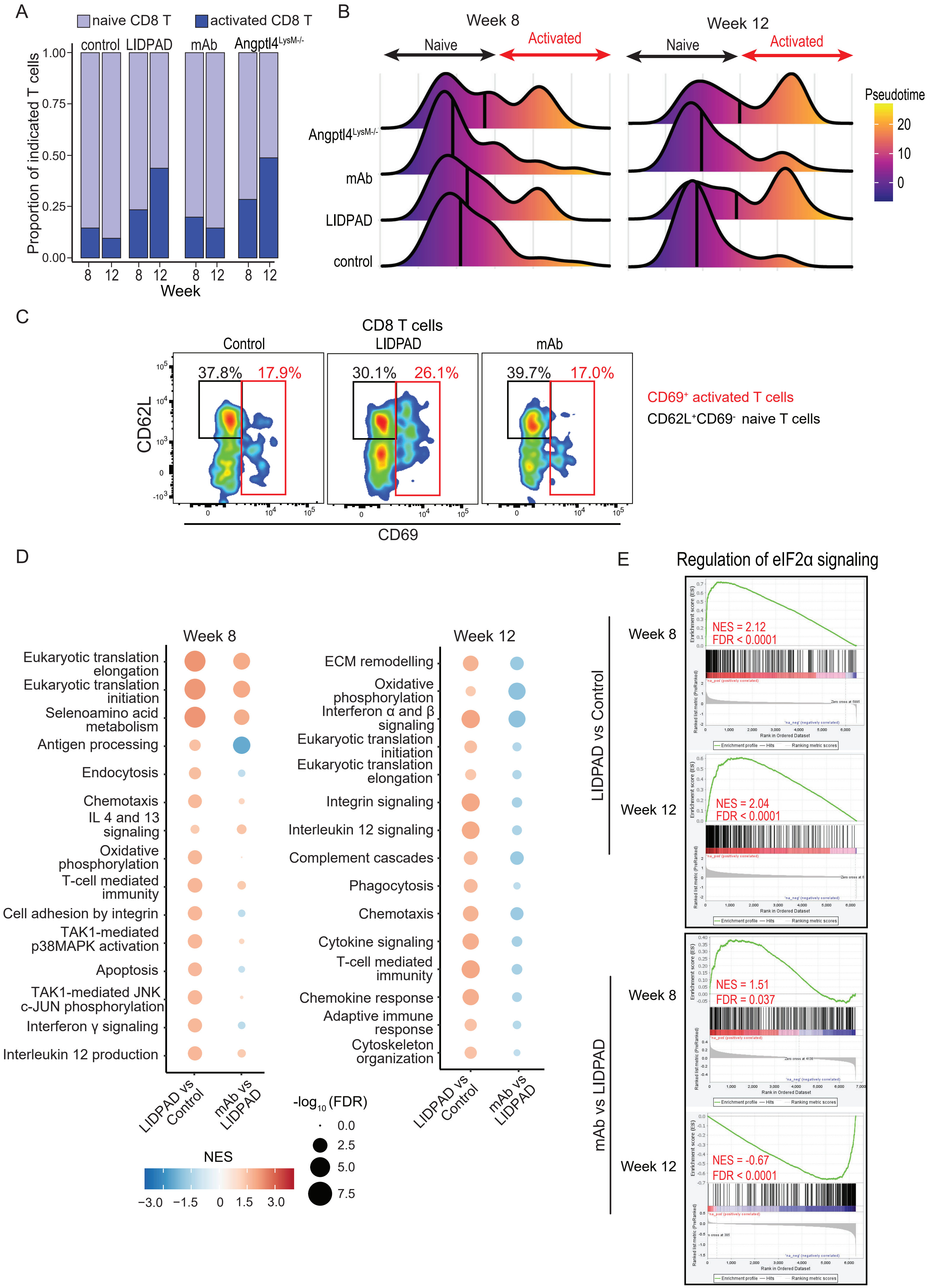
A
naive CD8 T
activated CD8 T
control
LIDPAD
mAb
Angptl4LysM-/-
Proportion of indicated T cells
1.00
0.75
0.50
0.25
0.00
8 12 8 12 8 12 8 12
Week
B
Week 8
Week 12
Naive
Activated
Angptl4LysM-/-
mAb
LIDPAD
control
Pseudotime
20
10
0
C
CD8 T cells
Control
LIDPAD
mAb
37.8%
17.9%
30.1%
26.1%
39.7%
17.0%
CD62L
CD69
CD69+ activated T cells
CD62L+CD69- naive T cells
D
Week 8
Eukaryotic translation elongation
Eukaryotic translation initiation
Selenoamino acid metabolism
Antigen processing
Endocytosis
Chemotaxis
IL 4 and 13 signaling
Oxidative phosphorylation
T-cell mediated immunity
Cell adhesion by integrin
TAK1-mediated p38MAPK activation
Apoptosis
TAK1-mediated JNK c-JUN phosphorylation
Interferon γ signaling
Interleukin 12 production
Week 12
ECM remodelling
Oxidative phosphorylation
Interferon α and β signaling
Eukaryotic translation initiation
Eukaryotic translation elongation
Integrin signaling
Interleukin 12 signaling
Complement cascades
Phagocytosis
Chemotaxis
Cytokine signaling
T-cell mediated immunity
Chemokine response
Adaptive immune response
Cytoskeleton organization
LIDPAD vs Control
mAb vs LIDPAD
-log10 (FDR)
0.0
2.5
5.0
7.5
NES
−3.0 −1.5 0 1.5 3.0
E
Regulation of eIF2α signaling
LIDPAD vs Control
Week 8
NES = 2.12
FDR < 0.0001
Week 12
NES = 2.04
FDR < 0.0001
mAb vs LIDPAD
Week 8
NES = 1.51
FDR = 0.037
Week 12
NES = -0.67
FDR < 0.0001

**Table S1. Nutritional Information on Diets (Experiment)**

| LIDPAD (Liver Disease Progression Aggravation Diet, Teklad Custom Diet, **1% cholesterol,** Envigo) | |
|---|---|
| Macronutrient Information | % kcal |
| **Protein** | 11.5 |
| **Carbohydrate** | 45.2 |
| **Fat** | 43.3 |

**Kcal/g: 4.6**

| Control (Teklad Custom Diet, Envigo) | |
|---|---|
| MacroNutrient Information | % kcal |
| **Protein** | 13.7 |
| **Carbohydrate** | 75.9 |
| **Fat** | 10.3 |

**Kcal/g: 3.6**

**Table S2. Relative abundance of intrahepatic immune cell subpopulations.**

| Subpopulations | Abundance (%) | | | | | | | |
|---|---|---|---|---|---|---|---|---|
| | Control | | LIDPAD | | Ab | | LysMCre-Angptl4$^{-/-}$ | |
| | Week 8 | Week 12 | Week 8 | Week 12 | Week 8 | Week 12 | Week 8 | Week 12 |
| Naive CD8 T | 14.35 | 12.28 | 7.14 | 9.77 | 7.79 | 13.56 | 17.88 | 2.17 |
| CD8 T cells | 2.58 | 1.36 | 2.23 | 7.27 | 1.97 | 2.35 | 7.11 | 2.07 |
| Naive CD4 T | 6.79 | 10.32 | 5.63 | 6.28 | 4.99 | 9.35 | 8.42 | 4.1 |
| CD4 T cells | 8.72 | 2.73 | 3.03 | 8.88 | 4.02 | 5.44 | 6.78 | 5.45 |
| dnT | 5.3 | 1.32 | 1.8 | 3.83 | 2.49 | 2.23 | 2.19 | 1.31 |
| T-innate-like | 0.68 | 0.64 | 1.02 | 0.88 | 1.62 | 0.59 | 0.44 | 0.14 |
| gdT | 2.22 | 1.36 | 2.97 | 6.49 | 3.24 | 3.11 | 3.01 | 2 |
| NKT | 12.62 | 3.37 | 4.69 | 5.8 | 6.65 | 4.72 | 5.96 | 1.59 |
| B cells | 21.05 | 52.35 | 56.26 | 37.03 | 49.08 | 42.65 | 21.82 | 61.75 |
| plasma cells | 0.41 | 0.32 | 0.35 | 0.1 | 0.26 | 0.34 | 0.11 | 0.79 |
| ILC | 2.45 | 0.96 | 2.72 | 2.34 | 3.94 | 2.06 | 2.62 | 1.48 |
| NK cells | 16.98 | 7.79 | 7.59 | 6.49 | 8.75 | 7.2 | 15.64 | 6 |
| Neutrophils | 0.23 | 0.24 | 0.55 | 0.36 | 1.4 | 0.27 | 1.59 | 4.14 |
| Monocytes | 1.18 | 0.4 | 2.19 | 0.88 | 1.66 | 0.82 | 2.08 | 2.31 |
| Kupffer cells | 0.08 | 0.08 | 0.22 | 0.19 | 0.13 | 0.56 | 0.11 | 0.66 |
| Mo-macs | 0.15 | 0.24 | 0.18 | 0.36 | 0.26 | 0.33 | 0.22 | 1.35 |
| cDC | 0.1 | 0.08 | 0.18 | 0.21 | 0.26 | 0.23 | 0.05 | 0.31 |
| pDC | 0.12 | 0.08 | 0.1 | 0.26 | 0.22 | 0.09 | 0.27 | 0.34 |
| 43 | 0.19 | 1.89 | 0.27 | 0.94 | 0.17 | 2.49 | 0.05 | 0.03 |
| 44 | 2.03 | 0.48 | 0.45 | 0.87 | 0.74 | 0.6 | 1.75 | 0.76 |
| 48 | 1.64 | 1.57 | 0.31 | 0.59 | 0.22 | 0.87 | 1.2 | 0.17 |
| 58 | 0.15 | 0.12 | 0.14 | 0.16 | 0.13 | 0.14 | 0.71 | 1.07 |

Total T cells and B cells include subpopulations highlighted in green and blue, respectively.

**Table S3. Relative abundance of T cell subpopulations**

| Subpopulation | Abundance (%) | | | | | | | |
|---|---|---|---|---|---|---|---|---|
| | Control | | LIDPAD | | Ab | | LysMCre-Angptl4$^{-/-}$ | |
| | Week 8 | Week 12 | Week 8 | Week 12 | Week 8 | Week 12 | Week 8 | Week 12 |
| Naive CD8 T | 26.95 | 36.78 | 25.03 | 19.85 | 23.77 | 32.79 | 34.53 | 11.54 |
| Memory CD8+ Tem | 3.62 | 3.12 | 5.9 | 10.3 | 4.94 | 4.26 | 8.13 | 5.13 |
| Memory CD8 T_c46 | 0.65 | 0.96 | 1.71 | 3.7 | 0.53 | 1.14 | 5.28 | 5.86 |
| Memory CD8+ Trm | 0.58 | 0 | 0.21 | 0.78 | 0.53 | 0.28 | 0.32 | 0 |
| Naive CD4 T | 12.75 | 30.89 | 19.75 | 12.76 | 15.22 | 22.61 | 16.26 | 21.79 |
| Treg_c13 | 10.58 | 2.52 | 3.7 | 5.71 | 3.74 | 4.57 | 5.6 | 10.07 |
| Treg_c27 | 2.93 | 3 | 3.43 | 5.04 | 4.27 | 2.04 | 4.22 | 9.89 |
| Th1 | 0.91 | 1.56 | 2.4 | 5.04 | 1.74 | 2.53 | 1.37 | 4.76 |
| Th17 | 1.96 | 1.08 | 1.1 | 2.26 | 2.54 | 4.02 | 1.9 | 4.21 |
| Naive dnT_c40 | 1.09 | 1.32 | 3.16 | 4.3 | 2.8 | 1.84 | 1.37 | 4.76 |
| Activated dnT_35 | 2.68 | 2.4 | 2.33 | 2.93 | 4.41 | 2.8 | 2.85 | 2.2 |
| Activated dnT_c51 | 6.19 | 0.24 | 0.82 | 0.56 | 0.4 | 0.76 | 0 | 0 |
| gdT | 3.3 | 3 | 4.87 | 4.69 | 7.08 | 4.92 | 3.38 | 2.01 |
| CCR6+ gdT | 0.87 | 1.08 | 5.56 | 8.5 | 2.8 | 2.6 | 2.43 | 8.61 |
| T-innate-like_c49 | 1.27 | 1.92 | 3.57 | 1.8 | 4.94 | 1.42 | 0.84 | 0.73 |
| NKT_c17 | 3.19 | 5.17 | 8.16 | 6.91 | 11.75 | 5.89 | 3.06 | 1.65 |
| NKT_c21 | 14.74 | 1.8 | 3.02 | 2.5 | 2.67 | 1.84 | 3.8 | 3.85 |
| NKT_c30 | 5.32 | 2.76 | 4.12 | 1.73 | 4.27 | 2.53 | 3.8 | 2.75 |
| Activated NKT_c55 | 0.43 | 0.36 | 1.17 | 0.63 | 1.6 | 1.18 | 0.84 | 0.18 |

**Table S4. Antibodies used for FACS**

| Antigen | Fluorophore | Cell type | Company | Catalog number |
|---|---|---|---|---|
| CD45 | FITC | Circulating leukocytes | BD | 553080 |
| CD45 | BV785 | Intrahepatic leukocytes | Biolegend | 103149 |
| CD11b | FITC | Myeloid cells | Miltenyi | 130-113-796 |
| NK1.1 | FITC | Natural killer cells | Biolegend | 156507 |
| Ly6G | FITC | Neutrophils | Miltenyi | 130-102-934 |
| CD19 | APC | B cells | Miltenyi | 130-123-791 |
| CD3 | BV421 | T cells | Biolegend | 100227 |
| CD4 | PE | CD4 T cells | Miltenyi | 130-116-509 |
| CD8b | BV711 | CD8 T cells | Biolegend | 126633 |
| CD62L | APC-Cy7 | Naïve T cells | Biolegend | 104427 |
| CD69 | BV510 | Activated T cells | Biolegend | 104531 |

**Table S5: List of kinase inhibitors used, molecular targets and concentration used in the drug screen.**

| Inhibitor | Target | Concentration (nM) |
|---|---|---|
| DMSO | NA | NA |
| Acalabrutinib | BTK | 20 |
| AMG-47a | Lck | 40 |
| AS703026 | MEK1/2 | 44 |
| AZD1208 | Pim1; Pim2; Pim3 | 80 |
| AZD-5438 | CDK1; CDK2; CDK9 | 80 |
| CC-401 HCl | JNK | 200 |
| CID755673 | PKD1; PKD2; PKD3 | 728 |
| CX-4945 | CK2 | 4 |
| Enzastaurin | PKCα; PKCβ; PKCγ; PKCε | 24 |
| GSK429286A | ROCK | 760 |
| H 89 diHydrochloride | PKA; S6K1 | 192 |
| IPI-145 | pan-PI3K | 200 |
| KN-62 | CaMK V ; CaMKII | 640 |
| KN-93 | CaMK | 1200 |
| Momelotinib | JAK1/2 | 80 |
| Palbociclib Isethionate | CDK4/6 | 64 |
| PD173955-Analog1 | c-Src | 40 |
| PF-562271 | FAK | 6 |
| PF-6260933 | MAP4K4 | 640 |
| Phenformin hydrochloride | AMPK | 40000 |
| R788 (Fostamatinib) Disodium | Syk | 164 |
| S-99 | ASK | 4 |
| SB202190 | MAPK (p38); p38αβ | 400 |
| SD169 | MAPK (p38) | 16 |
| SNS-314 | Aurora A/B/C | 124 |
| Tofacitinib | JAK3 | 364 |

**Supplemental Figure Legends**

**Figure S1: Metabolic parameters of control and LIDPAD mice.**

**A.** Change in weight (left) of mice fed the control diet and LIDPAD and the number of animals (left) measured at each time point for 48 weeks**.**

**B.** IGTT curves of LIDPAD- and control-fed mice from weeks 1 to 48.

n = 7-10 per group. Data are expressed as the means ± SEMs. ****p<0.0001, ***p<0.001, **p<0.01, *p<0.05 (unpaired t test, ANOVA Welch's t test or ANCOVA test when appropriate, followed by post hoc comparisons).
